# Translational control of *MPS1* links protein synthesis with the initiation of cell division and spindle pole body duplication in *Saccharomyces cerevisiae*

**DOI:** 10.1101/2020.12.29.424704

**Authors:** Heidi M. Blank, Annabel Alonso, Amy S. Fabritius, Mark Winey, Michael Polymenis

## Abstract

Protein synthesis underpins cell growth and controls when cells commit to a new round of cell division at a point in late G1 of the cell cycle called Start. Passage through Start also coincides with the duplication of the microtubule-organizing centers, the yeast spindle pole bodies, which will form the two poles of the mitotic spindle that segregates the chromosomes in mitosis. The conserved Mps1p kinase governs the duplication of the spindle pole body in *Saccharomyces cerevisiae*. Here, we show that the *MPS1* transcript has a short upstream open reading frame that represses the synthesis of Mps1p. Mutating the *MPS1* uORF makes the cells smaller, accelerates the appearance of Mps1p in late G1, and promotes completion of Start. Monitoring the spindle pole body in the cell cycle using structured illumination microscopy revealed that mutating the *MPS1* uORF enabled cells to duplicate their spindle pole body earlier at a smaller cell size. The accelerated Start of *MPS1* uORF mutants depends on the G1 cyclin Cln3p and the transcriptional repressor Whi5p but not on the Cln1,2p G1 cyclins. These results identify growth inputs in mechanisms that control duplication of the microtubule-organizing center and implicate these processes in the coupling of cell growth with division.

## INTRODUCTION

Basic cellular parameters, such as cell size and macromolecular composition, do not change much throughout successive cell divisions, and cell growth and division appear balanced (Polymenis 2022). The rate at which cells can divide usually depends not on how fast they can replicate their genome but on how fast they can synthesize everything else that goes into a newborn cell (Johnston *et al*. 1977). Newborn budding yeast daughter cells actively control when they begin replicating their genome in response to their growth status. They stay in the G1 phase of the cell cycle until they reach a characteristic threshold of cell growth known as critical size (Hartwell and Unger 1977; Di Talia *et al*. 2007). The protein synthesis rate is thought to underpin overall cell growth and determine when cells will reach their critical size and commit to a new round of DNA replication at Start (Polymenis and Aramayo). Translational control of *CLN3*, encoding a G1 cyclin, alters the timing of Start in different nutrients (Polymenis and Schmidt 1997; Blank *et al*. 2018), and synthesis of Cln3p in G1 may respond to a critical rate of protein synthesis (Litsios *et al*. 2019). However, other molecular targets that could impose the protein synthesis requirements for cell division initiation have remained elusive.

The spindle pole body (SPB) of yeast is a large protein organelle serving the same role as the centrosome in animals. The two SPBs in mitosis form the poles from which all the microtubules of the mitotic spindle originate (Jaspersen and Winey 2004). The SPB is a stack of three major protein layers or plaques embedded in the nuclear envelope. Besides the chromosomes, the SPB is the only large macromolecular complex present in a single copy early in the G1 phase of the cell cycle. When cells pass through Start, a protein extension of the middle plaque called the half-bridge nucleates the new SPB formation, maturing and separating from the old SPB. From classical experiments involving loss-of-function mutants, it is thought that passage through Start is required for SPB duplication and not vice versa (Hartwell *et al*. 1974; Pringle 1981).

Mps1p is a conserved protein serine, threonine, and tyrosine kinase required for SPB duplication (Winey *et al*. 1991; Lauzé *et al*. 1995; Liu and Winey 2012). Mps1p also regulates various other mitotic events, most prominently bipolar chromosome attachment and the spindle checkpoint (Liu and Winey 2012). Post-translational mechanisms control Mps1p’s levels (through proteolysis), activity (through phosphorylation by several kinases, including by Mps1p itself), and localization (at least in part through phosphorylation and localized degradation) in the cell (Liu and Winey 2012). Mps1p abundance is very low, at ∼500 molecules/cell in asynchronous cells (Ho *et al*. 2018). Mps1p levels peak in metaphase, consistent with Mps1p’s numerous mitotic roles, but then the anaphase-promoting complex destroys Mps1p for a timely mitotic exit (Palframan *et al*. 2006). Over-expression of Mps1p arrests cells in mitosis (Hardwick *et al*. 1996). How Mps1p levels are controlled in G1 is unknown.

We previously identified *MPS1* as one of a few mRNAs whose translational efficiency changes in the cell cycle (Blank *et al*. 2017). Here, we identify a short upstream open reading frame (uORF) in *MPS1*. Mutating the *MPS1* uORF caused an earlier appearance of Mps1p in G1 and accelerated SPB duplication. It also reduced the critical size for Start completion in a Cln3p-dependent manner. These results reveal direct growth and protein synthesis inputs in the Mps1p-controlled SPB cycle. They also show that SPB-related processes can actively determine when cells commit to a new round of cell division at Start.

## MATERIALS AND METHODS

### Key resources table

Where known, the Research Resource Identifiers (RRIDs) are shown.

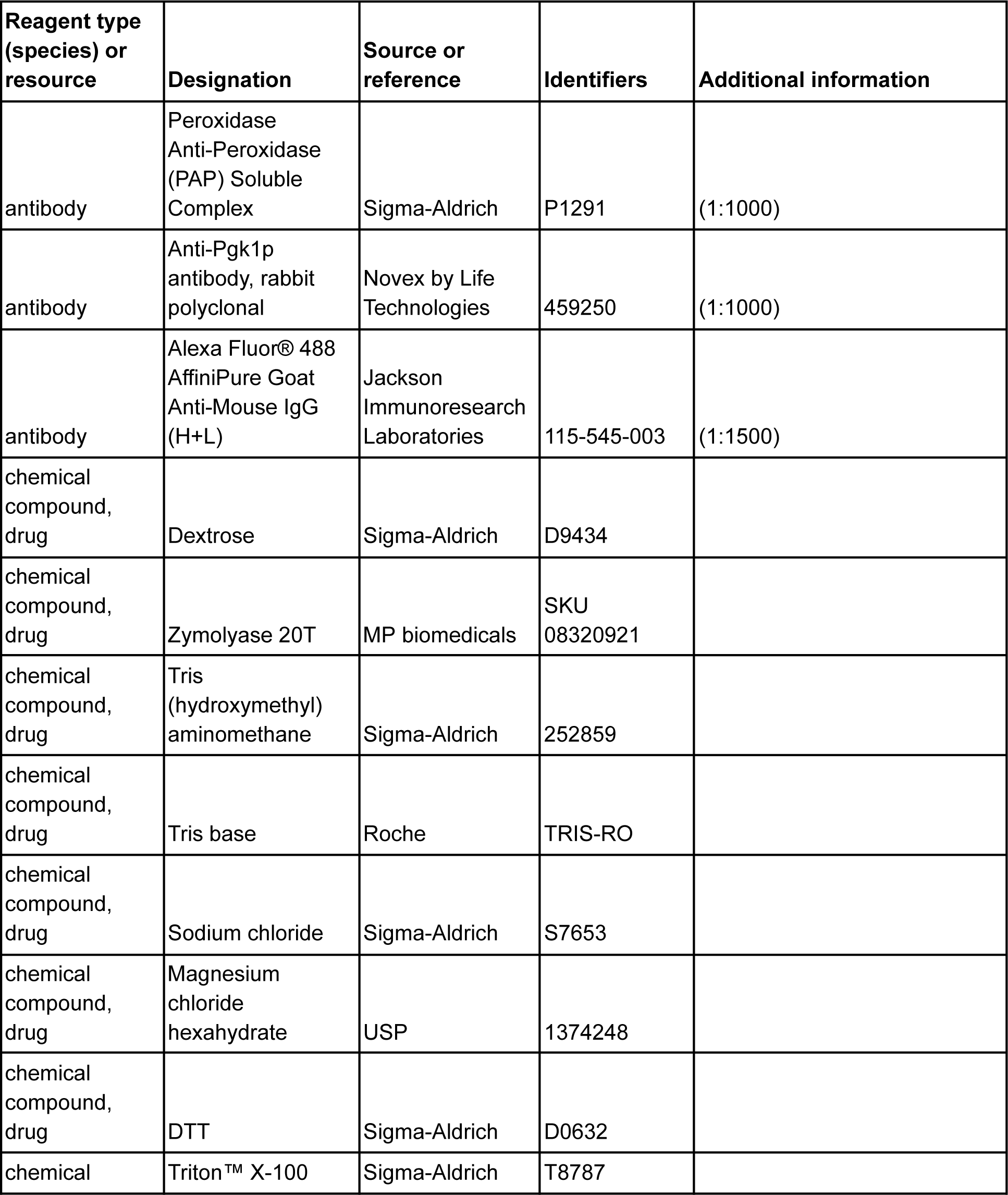
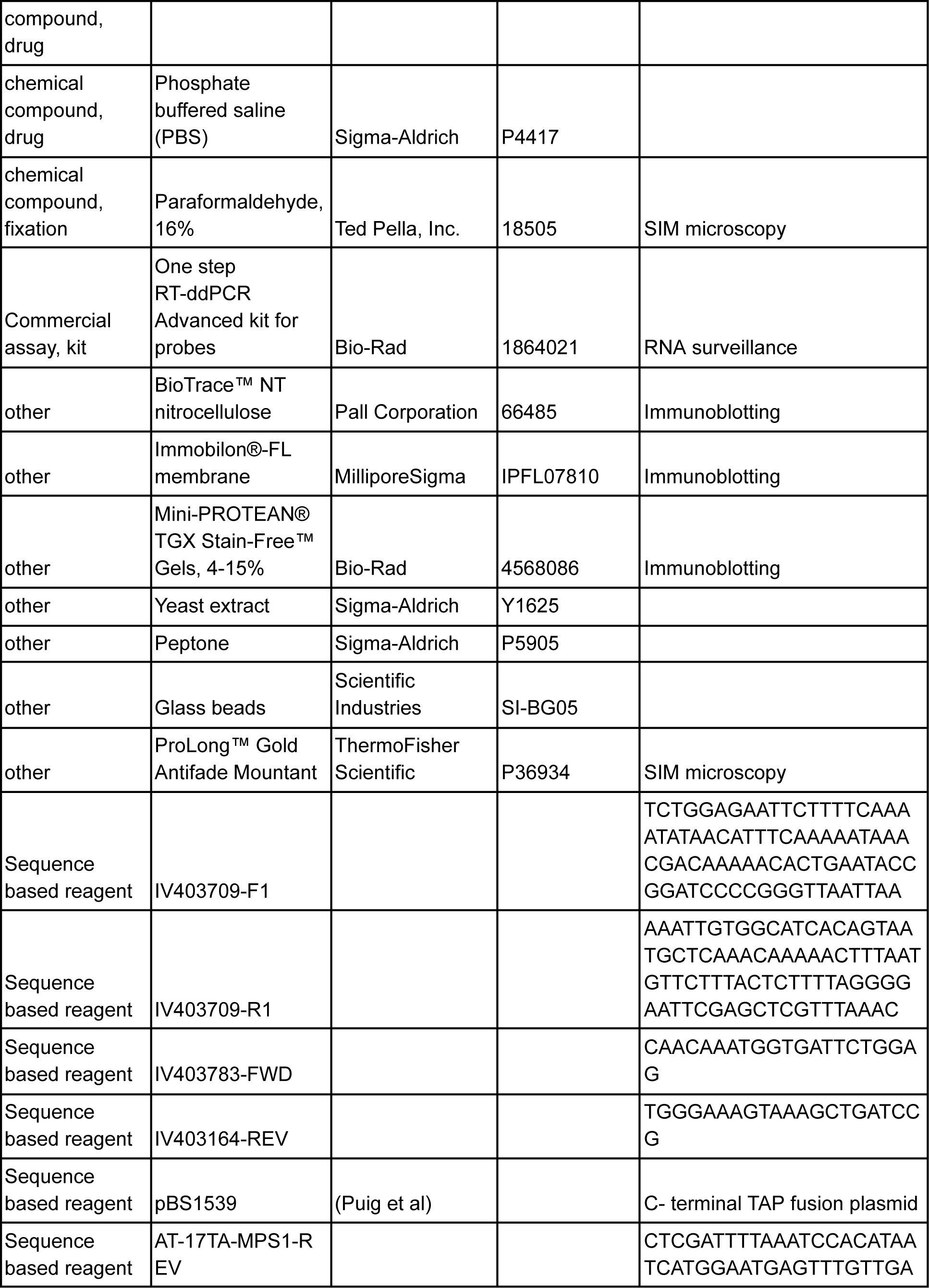
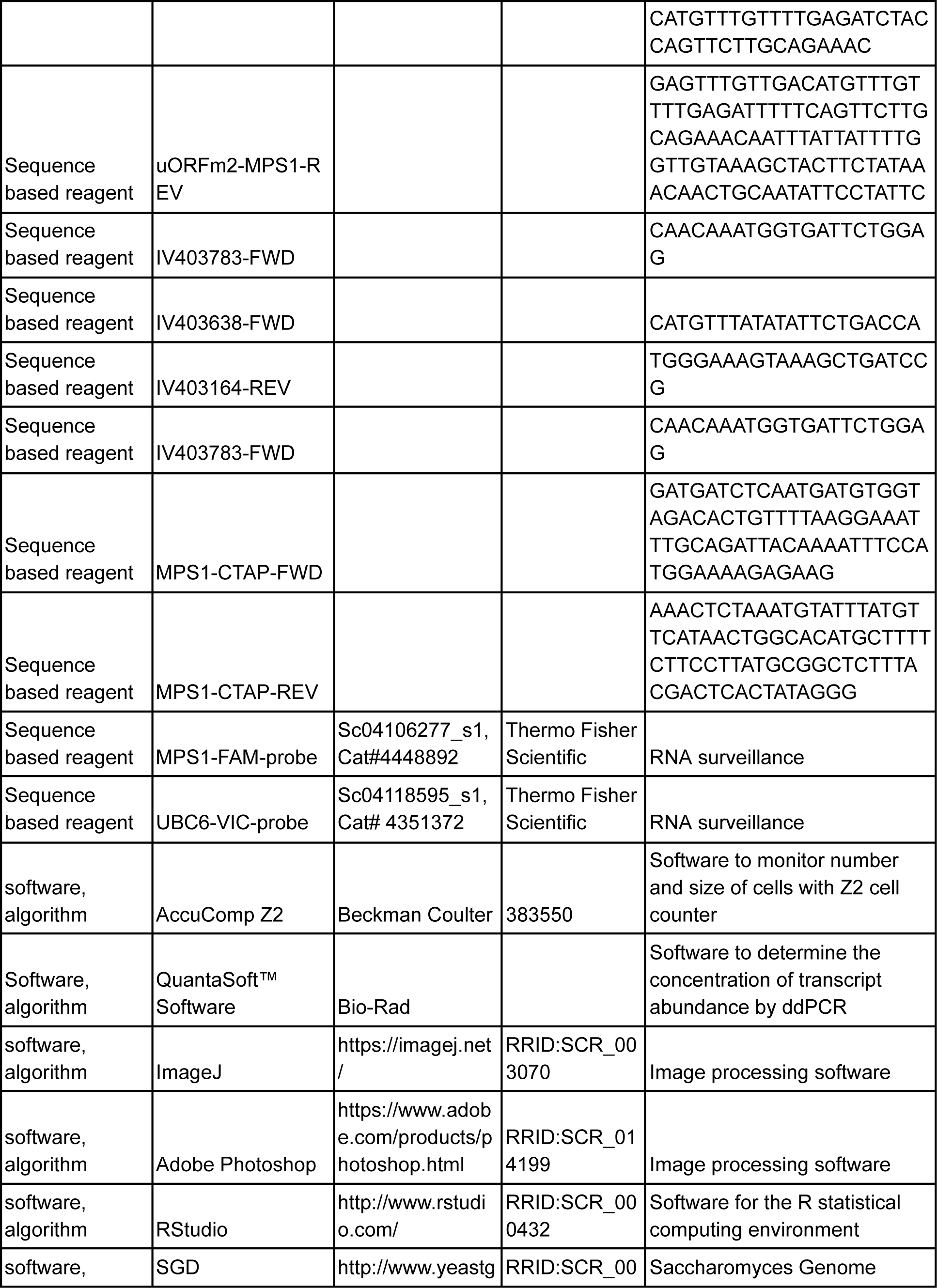
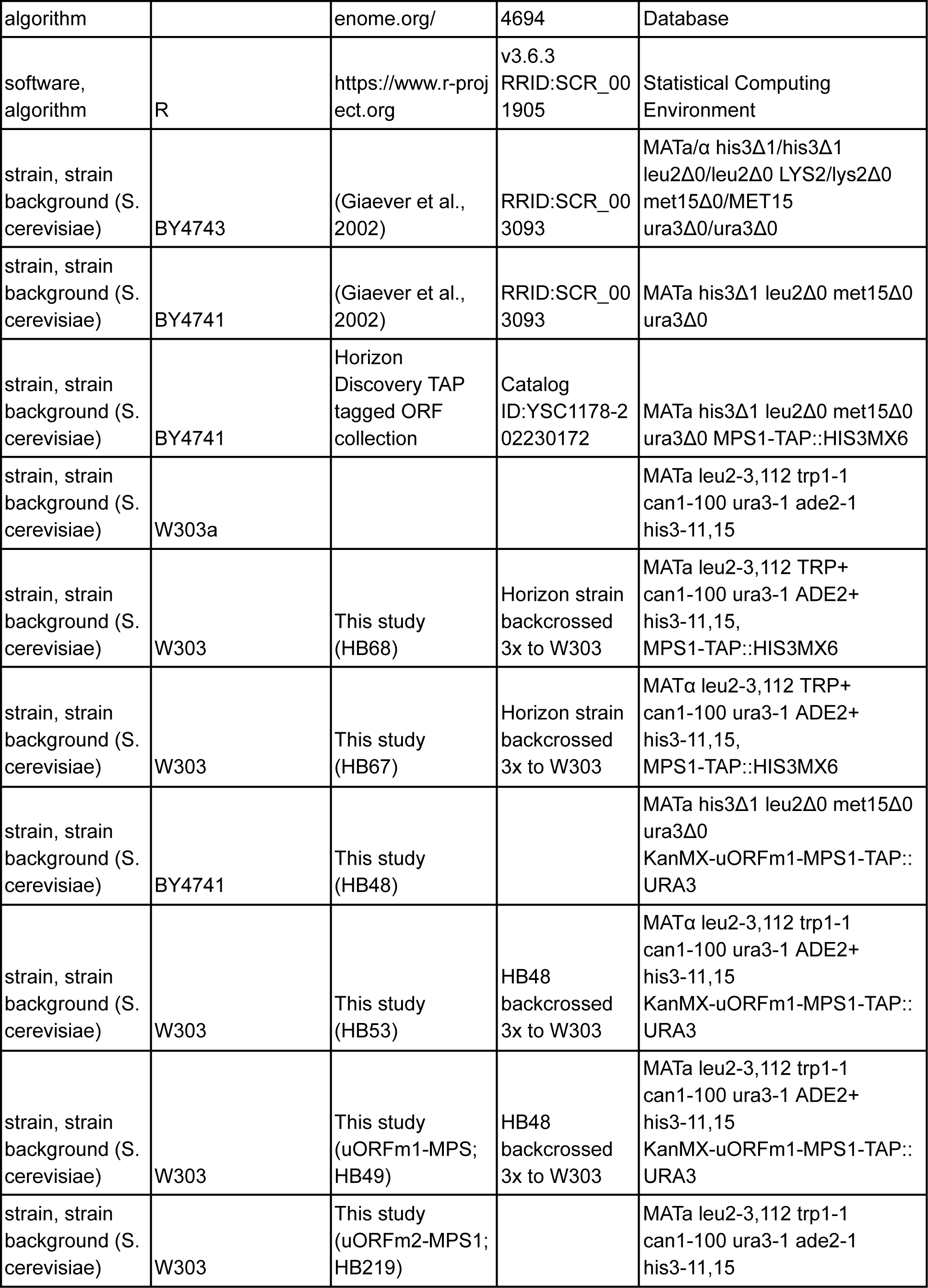
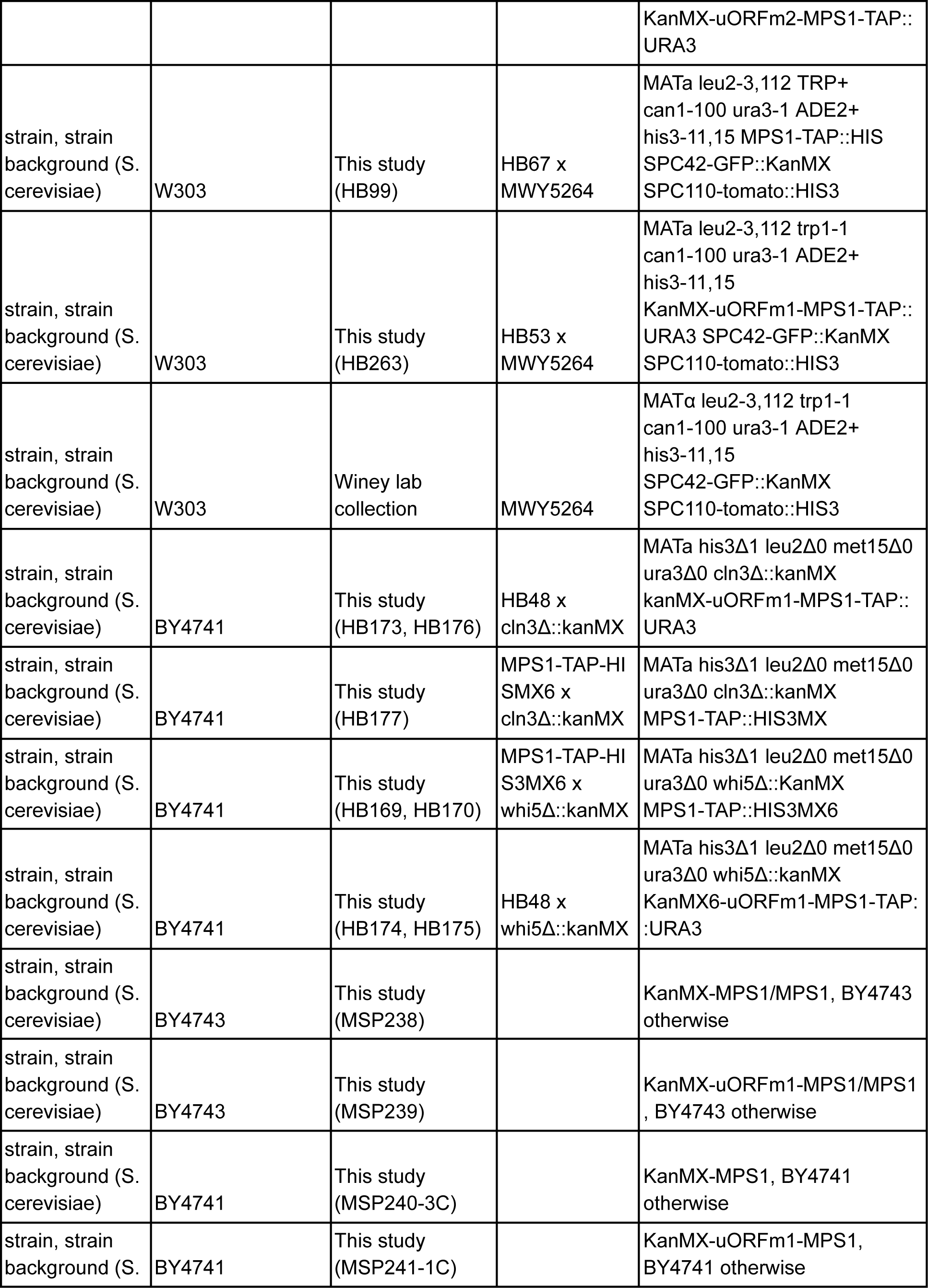
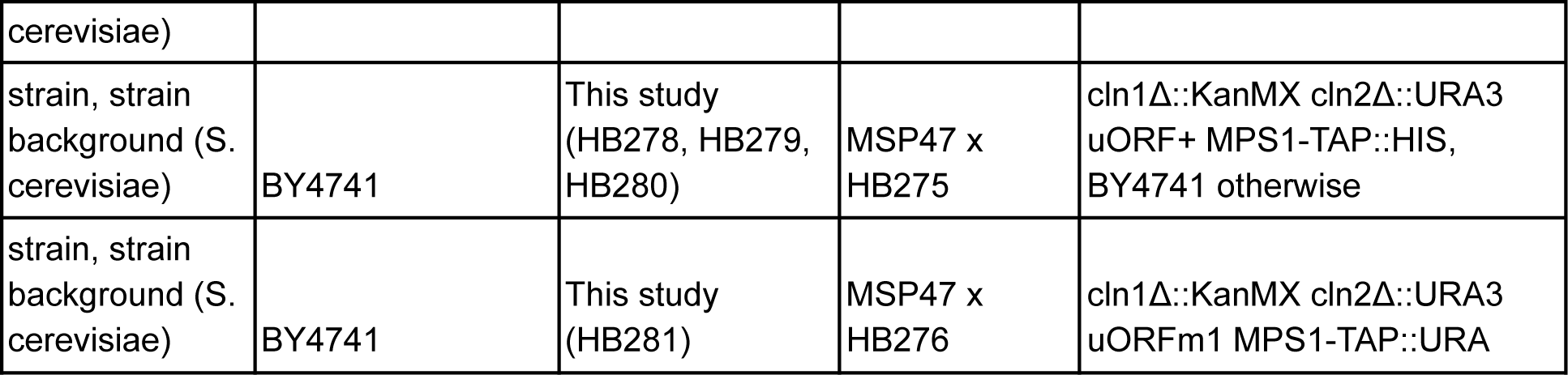

### Strains and media

All the strains used in this study are shown in the Key Resources Table above. Unless noted otherwise, the cells were cultivated in the standard, rich, undefined medium YPD (1% ^w^/_v_ yeast extract, 2% ^w^/_v_ peptone, 2% ^w^/_v_ dextrose), at 30°C (Kaiser *et al*. 1994).

To modify the *MPS1*-5’ leader, we first inserted a *KanMX6* cassette upstream, at position ChrIV:403709 with the PCR-mediated methodology of (Longtine *et al*. 1998), using primers IV403709-F1 and IV403709-R1 (see Key Resources Table). The PCR product was introduced into strain BY4743, generating the heterozygous strain MSP238, which was sporulated and dissected to generate the haploid MATɑ (MSP240-3A) and MATa (MSP240-3C) segregants. The *KanMX6* insertion was verified by PCR, using primers IV403783-FWD and IV403164-REV.

To introduce the uORFm1 mutation, we used primers IV403783-FWD and AT-17TA-MPS1-REV in a PCR reaction with genomic DNA from MSP240-3A. The PCR product was then used to transform strain BY4743, generating strain MSP239. The introduction of the uORFm1 was initially verified by restriction digestion since the uORFm1 introduces a BglII restriction site. Strain MSP239 was then sporulated and dissected to generate the haploid uORFm1 haploid MATa (MSP240-1C) segregant. The uORFm1 mutation was verified by sequencing with primers IV403638-FWD and IV403164-REV.

To insert a C-terminal TAP tag at the *MPS1* locus, we amplified the TAP-tagged sequences from plasmid pBS1539 (Puig *et al*. 2001) with primers MPS1-CTAP-FWD and MPS1-CTAP-REV and used the PCR product to transform the above generated *uORF^+^-MPS1* and *uORFm1-MPS1* haploid strains. These BY4741-based strains were then crossed with W303ɑ three times to generate the equivalent W303-based strains.

For constructing the uORFm2-MPS1-TAP strain, primers uORFm2-MPS1-REV and IV403783-FWD were used in a PCR reaction, using as template genomic DNA of a uORFm1-MPS1-TAP strain. The generated PCR product contained the KanMX cassette and introduced the point mutations at the uORF of *MPS1* shown in Figure 1. This PCR product was transformed into W303a cells, and potential candidates were sequenced with primers IV403638-FWD and IV403164-REV to confirm the presence of the uORF mutations. Verified uORFm2 strains were then transformed again with the PCR product of primer pair MPS1-CTAP-FWD and MPS1-CTAP-REV with DNA extracted from the uORFm1 strain to insert the TAP protein tag. All transformed strains were backcrossed once.

**Figure 1.**
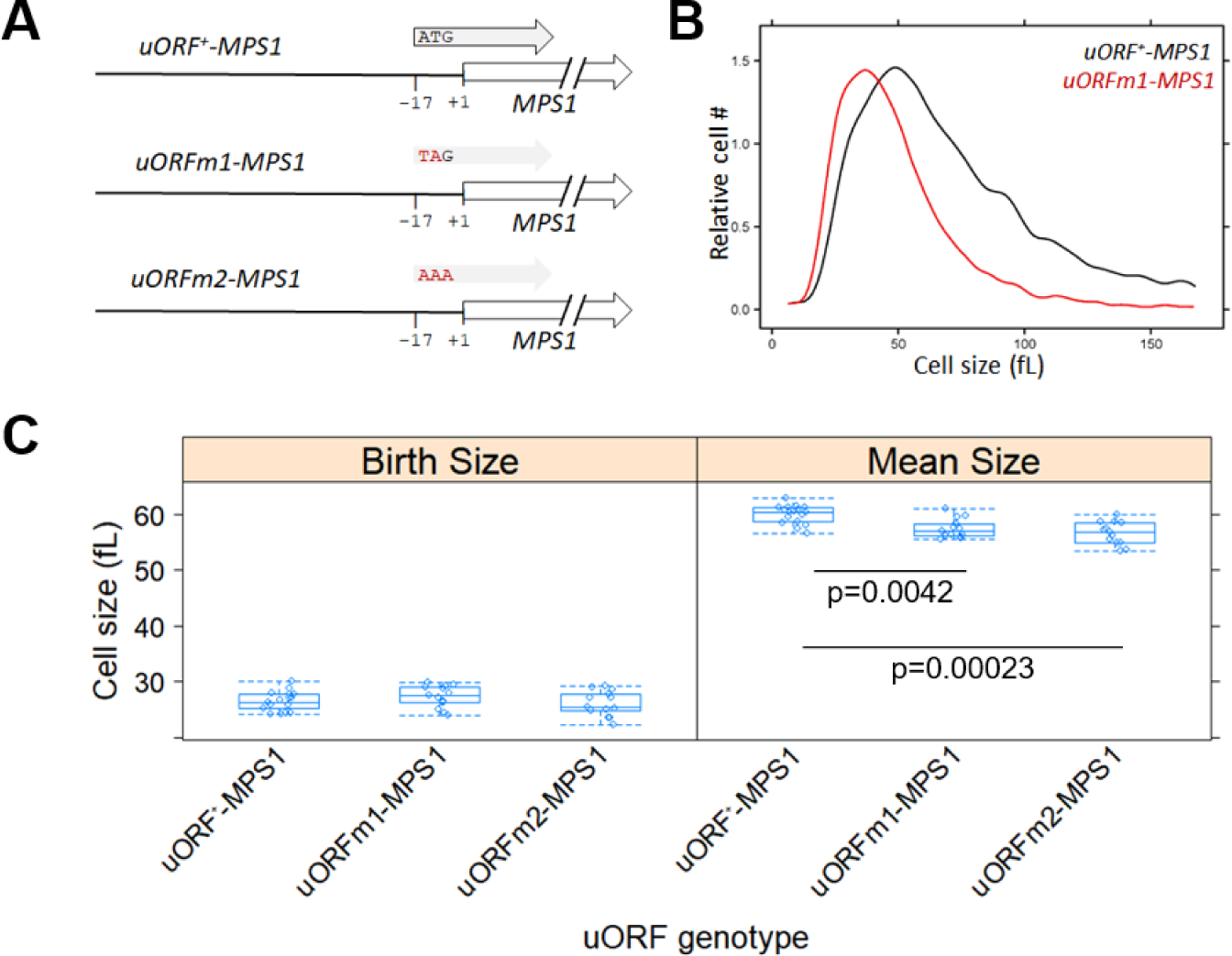
Mutating a conserved uORF in *MPS1* reduces cell size. **A**, Schematic of the *MPS1* locus in the strains used in this study to test the role of the *MPS1* uORF. **B**, Cell size histograms (size in fL, x-axis; number of cells, y-axis) for the indicated strains (*uORF^+^-MPS1* [strain MSP240-3C] and *uORFm1-MPS1* [strain MSP241-1C]; see Materials and Methods) in otherwise wild type cells, cultured in rich medium (YPD), proliferating asynchronously and exponentially. **C**, Box-plots showing the mean and birth size of populations of cells from multiple independent measurements from the indicated strains (all in the W303 background, carrying TAP-tagged *MPS1* alleles; see Materials and Methods). The associated p-values were from the nonparametric Kruskal-Wallis and the posthoc Nemenyi tests (calculated with the PMCMR R language package) between the samples shown. The values used to generate the graphs in C are in supplementary File1/sheet ‘fig1c’.

We constructed the fluorescently tagged strains for SIM microscopy by crossing strain MWY5624 with the strains carrying the appropriate TAP-tagged *uORF+-MPS1* or *uORFm1-MPS1* allele. Since these crosses involved multiple genes tagged with the same marker, the obtained tetrads were analyzed carefully to pick non-parental ditypes of the double genetic markers. Strains were further confirmed by immunoblotting for Mps1p-TAP, and both fluorescent markers were confirmed microscopically.

### Sample-size and replicates

For sample-size estimation, no explicit power analysis was used. All the replicates in every experiment shown were biological ones from independent cultures. A minimum of three biological replicates was analyzed in each case, as indicated in each corresponding figure’s legends. Where three replicates were the minimum number of samples analyzed, the robust bootstrap ANOVA was used to compare different populations via the t1waybt function, and the posthoc tests via the mcppb20 function of the WRS2 R language package (Wilcox 2011; Mair and Wilcox 2020). We used nonparametric statistical methods for measurements where at least four independent replicates were analyzed, as indicated in each case. No data or outliers were excluded from any analysis.

### Centrifugal elutriation and cell size measurements

For each elutriation experiment in YPD (1% ^w^/_v_ yeast extract, 2% ^w^/_v_ peptone, 2% ^w^/_v_ dextrose, plus 0.02 g/L of adenine for strains of W303 background), a 350 mL culture was allowed to reach a density of 1-5E+07 cells/mL. Cells were loaded at a 35 mL/min pump speed onto a large elutriator chamber (40 mL), spinning at 3,200 rpm (Beckman J-6M/E centrifuge). The centrifuge chamber was set at 4°C during all centrifugation steps until approximately 10 min before cell harvesting, when the temperature was increased to avoid the refrigeration from turning on and disrupting the cell gradient in the chamber. For typical time-series experiments with synchronous elutriated cultures (e.g., see Figure 2A), we isolated the first, early G1 daughter fraction and monitored it at regular intervals afterward as it progressed in the cell cycle, as we have described elsewhere (Hoose *et al*. 2012; Soma *et al*. 2014).

**Figure 2.**
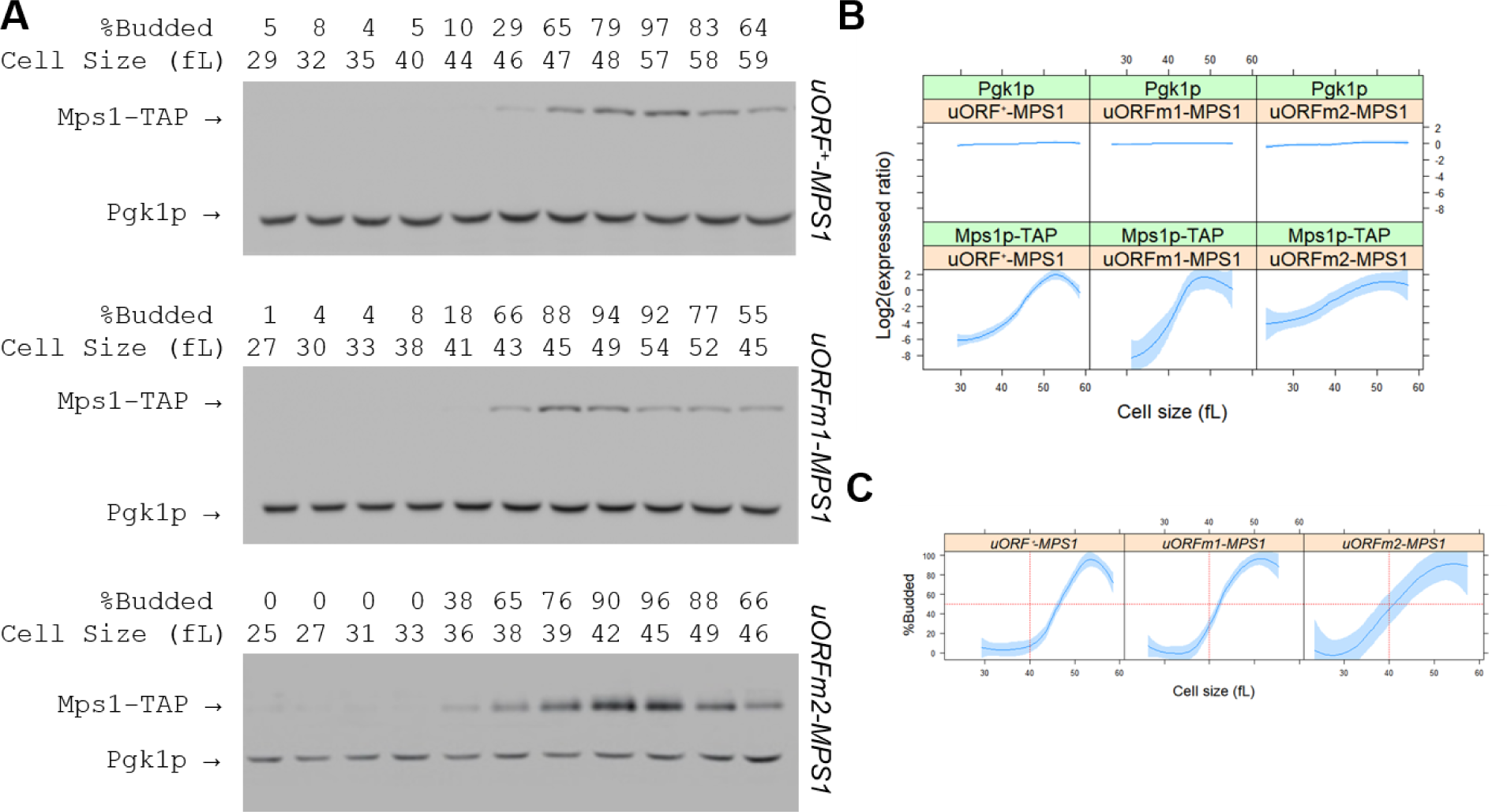
Mutating the *MPS1* uORF accelerates the appearance of Mps1p in the cell cycle and promotes Start. The abundance of Mps1p-TAP was monitored in wild type and *uORFm-MPS1* cells, constructed as described in Materials and Methods. Samples were collected by elutriation in a rich, undefined medium (YPD) and allowed to progress synchronously in the cell cycle. Experiment-matched loading controls (measuring Pgk1p abundance) were also quantified and shown in parallel. **A**, Representative immunoblots, along with the percentage of budded cells (%B) and the cell size (in fL), for each of the samples. **B**, From at least three independent experiments in each case, the Mps1p-TAP and Pgk1p signal intensities were quantified as described in Materials and Methods. The Log2(expressed ratios) values are on the y-axis, and cell size values are on the x-axis. Loess curves and the standard errors at a 0.95 level are shown. All the immunoblots for this figure are in supplementary File2, while the values used to generate the graphs are in supplementary File1/sheet ‘fig2b’ and sheet ‘fig2c’.

### Protein surveillance

Protein extracts for immunoblots were made with the sodium hydroxide extraction method described previously (Amberg *et al*. 2006). The extracts from synchronous culture samples were run on 10% Tris-Glycine SDS–PAGE gels and transferred to a nitrocellulose membrane. For detection of proteins of interest on immunoblots, we used an anti-Pgk1p antibody to detect Pgk1p and the peroxidase-anti-peroxidase (PAP) soluble complex to detect TAP-tagged proteins. Chemiluminescence reagents were from Thermo Scientific and used at the dilutions recommended by the manufacturer for detection of the TAP-tagged Mps1p. A secondary antibody was used to detect Pgk1p (Jackson Immunoresearch Laboratories). Detection of the proteins was digitally acquired using an Amersham™ Imager 600 (GE Healthcare).

The extracts from batch cultures were prepared the same as above but instead ran on Mini-PROTEAN® TGX Stain-Free™ Gels and transferred to a low-fluorescence PVDF membrane. Protein detection was as above, but using the stain-free gel technology, we could acquire a total protein image of the blot to be used as an additional loading control along with Pgk1p. The manufacturer’s instructions were followed to activate the gel, except using an Amersham™ Imager (GE Healthcare) for 3 minutes to activate the gel after completion of electrophoresis. Following transfer to an Immobilon®-FL membrane, the total protein was detected using the UV channel of the Amersham™ Imager 600. Images were processed with the “Subtract Background” tool, and band intensities quantified using the “Measure” tool to obtain a mean intensity for each band, using the ImageJ software package. The area measured was kept constant for a particular sample series for each blot analyzed.

To calculate the Log2(expressed ratios) in cell cycle series (e.g., see Figure 2), for any one protein of interest (Mps1p-TAP or Pgk1p), each signal value was divided by the average signal across the entire cell cycle series for that protein. These ratios were then Log2-transformed and reported on the y-axis.

### Transcript abundance using digital droplet PCR (ddPCR)

For RNA surveillance, RNA extracts were made as described previously (Blank *et al*. 2020). Briefly, RNA was extracted using the hot acidic phenol method. Cell pellets were resuspended in 0.4 ml of TES buffer (10 mM Tris, pH 7.5; 10 mM EDTA, and 0.5% SDS) with 0.1 ml of glass beads. Then, 0.4 ml of acid phenol, pH 4.5, was added, and the samples were heated at 65°C for 0.5 h with occasional vortexing for 10 s each time. The samples were then centrifuged for 5 m at 13,000 g. The aqueous layer was transferred to a new tube containing 1 ml of cold 100% ethanol with 40 μl of 3 M sodium acetate and incubated overnight at 4°C. The next day, the samples were centrifuged at 13,000 g for 20 min at 4°C, washed with 80% ethanol, and centrifuged again for 5 min at 13,000 g. The pellets were resuspended in 25 μl water. The amount of total RNA in each sample was measured with a ThermoFisher NanoDrop™ spectrophotometer. For the quantification of transcript abundance, 1 ng of total RNA was used for each sample.

The ddPCR reaction mixture was prepared by following the manufacturer’s protocol (One-Step RT-ddPCR Advanced Kit for Probes), using the TaqMan® hydrolysis probes labeled with FAM-MBP for MPS1 and VIC-MBP for UBC6 reporter fluorophores. The mixture was kept on ice throughout the whole experiment. Once the reaction mixture was prepared, the samples were placed into a droplet generator (QX200™ AutoDG™ Droplet Digital™ PCR System), which uses specially developed reagents and microfluidics to partition each sample into 20,000 nl-sized droplets. Once the droplets were generated, the samples were transferred to a thermal cycler (C1000 Touch™ Thermal cycler) for PCR. Following the PCR, the plate containing the droplets was placed in a droplet reader (Bio-Rad, QX200 Droplet reader). he droplet reader’s autosampler picks up droplets from each of the wells of the PCR plate, and fluorescence is measured for individual droplets. The abundance of the transcripts was obtained using the QuantaSoft™ Software. Transcript levels of *MPS1* were normalized against the corresponding transcript levels of *UBC6*.

### Cell fixation for SIM microscopy

Cells containing the fluorescent markers for SPB analysis were elutriated as described earlier. For each time point, 1 mL of cell culture was collected, pelleted, and washed in 1mL of PBS. Cells were then spun at 3,000rpm for 3 m in a microcentrifuge and resuspended in 1mL of 4% paraformaldehyde (Ted Pella, Inc.) in a 133mM sucrose in PBS solution. Fixation was for 15 min at room temperature, followed by two more washes in PBS. Cells were stored at 4°C until they were analyzed by microscopy.

### SIM microscopy

Structured Illumination Microscopy analysis was performed on fixed cells, which had been mounted onto glass slides using ProLong Gold Antifade mountant (ThermoFisher Scientific). Mounted cells were stored in the dark at room temperature for 48 h prior to SIM analysis. Images of Z-series (step size of 0.2 μm) of cells were collected using the Nikon Structured Illumination Super-Resolution Microscope fitted with a 100×1.46 NA objective. Images were captured in the 3D-SIM mode using NIS-Elements and SIM processing was performed using the SIM module in the Nikon software package. Image projections were made in Image-J software (National Institutes of Health).

### In vitro phosphorylation assays

All methods for protein purification and kinase reactions have been described previously (Venta *et al*. 2020). Ndc80p (1-257 aa) and Sic1p (1-215 aa) were expressed and purified as 6xHis fusions. Sic1p-AP had all known Cdk sites mutated to Ala. Whi5p, Cdc31p, and Spc29p were expressed and purified as GST fusions of the full-length protein in each case. In Figure S5A, the enzyme concentration was 2 nM, and the substrate protein was 1 μM. In the experiments in Figure S5B, Mps1p was present at 4.2 nM, and the protein substrates were at 350 nM. After SDS-PAGE, gels were vacuum-dried. The BAS-IP MS 2040 E screen, Biomolecular Imager Typhoon 5, and ImageQuant TL software (all from Amersham) were used for autoradiography.

## RESULTS AND DISCUSSION

### Mutating a conserved uORF in *MPS1* reduces cell size

To test the role of translational control in Mps1p synthesis, we looked at possible *cis*-elements in the *MPS1* mRNA that could mediate that control. The 5’-leader of *MPS1* is 133 nt-long (Poch *et al*. 1994) and has a uORF (Figure S1, top). The uORF initiates 17 nt upstream but overlaps and terminates past the main *MPS1* start codon (Figure S1). The uORF could repress the translation of *MPS1* because ribosomes that synthesize the uORF-encoded peptide will bypass the *MPS1* start codon. A uORF at the same position, albeit of varying length and sequence, is present in other *Saccharomyces* genomes (Figure S1). Based on aggregate ribosome profiling experiments from the literature, the uORF sequences are bound by ribosomes (Michel *et al*. 2014), suggesting that the uORF is translated.

To test the role of the *MPS1* uORF, we mutated its start codon in two ways (Figure 1A). The 5’-leader may also be structured (Figure S2A). The uORF mutations we introduced may (*uORFm1-MPS1*) or not (*uORFm2-MPS1*) alter the predicted 5’-leader structure (Figure S2A). The two different mutants provide independent means to test the role of the *MPS1* uORF. We introduced these mutations in their native context in the chromosome, with the endogenous *MPS1* promoter driving the *MPS1* mRNA expression (see Materials and Methods). We also introduced a C-terminal TAP tag to visualize Mps1p-TAP on immunoblots (see Materials and Methods).

While both *MPS1* uORF mutants proliferated at the same rate as their *uORF^+^* counterpart, they had a small overall cell size, regardless of strain background (BY4741 in Figure 1B; W303 - Figure 1C). The cells were born at the right size (Figure 1C, left panel), but the mean size of asynchronously proliferating cells was about 10-15% smaller (Figure 1C, right panel; p<0.005 based on the non-parametric Kruskal-Wallis and the posthoc Nemenyi tests). The uORF mutations did not change the steady-state levels of *MPS1* mRNA or Mps1-TAP protein in asynchronous cells (Figure S2B). This is not surprising due to the other known mechanisms regulating Mps1p abundance in the cell cycle, as described in the next section. However, the smaller mean cell size of asynchronous cultures of the *MPS1* uORF mutants was suggestive of cell cycle-dependent phenotypes, which we examined next.

### Mutating the *MPS1* uORF accelerates the appearance of Mps1p in the cell cycle and promotes Start

Smaller cell size is often associated with accelerated completion of Start, as shown initially for stabilized, gain-of-function mutants of the G1 cyclin Cln3p (Sudbery *et al*. 1980; Cross 1988). To measure the timing of Start in the *MPS1* uORF mutants, we obtained highly synchronous cultures by centrifugal elutriation, which maintains as much as possible the coordination of cell growth with division (Creanor and Mitchison 1979; Aramayo and Polymenis 2017). The abundance of Mps1p in elutriated daughter cells has not been measured in past studies. We found that Mps1p-TAP is absent in G1 and begins accumulating at the G1/S transition when cells begin budding (Figure 2A, top panel). This is expected because Mps1p is a substrate of the anaphase-promoting complex (APC) (Palframan *et al*. 2006), and the APC is active in G1. Mps1p-TAP levels peak late in the cell cycle, in mitosis, when the percentage of budded cells is the highest (>90%), and then Mps1 levels drop again as cells exit mitosis (Figure 2A, B). In both independently constructed *MPS1* uORF mutants, Mps1p-TAP abundance was also periodic, absent in G1, and peaking late in the cell cycle (Figure 2A, B). However, Mps1p-TAP began appearing in smaller cells (∼5 fL smaller) in the *MPS1* uORF mutants than wild-type cells (Figure 2A, B). Hence, although mutating the *MPS1* uORF does not change the steady-state levels of Mps1p in asynchronous cells (Figure S2B), it changes the kinetics of the Mps1p appearance in the cell cycle (Figure 2).

Budding yeast daughter cells commit to a new round of DNA replication when they reach a critical size threshold at Start. Since budding coincides with the initiation of DNA replication in yeast (Hartwell *et al*. 1974), from such synchronous elutriated cultures, the critical size is commonly defined as the size at which half the cells are budded. Remarkably, the critical size of *MPS1* uORF mutants was ∼40 fL, while that of wild type cells was >45 fL (Figure 2C; e.g., the critical size is reduced by 9% (Wilcoxon rank-sum test; p-value = 0.0005828) in the *uORFm1-MPS1* mutant). We conclude that Start was accelerated in the *MPS1* uORF mutants.

### Reducing the translational efficiency of *MPS1* accelerates SPB duplication in synchronous, elutriated cells

Since Mps1p plays multiple roles in SPB duplication, we examined the effect of the *uORFm1-MPS1* mutation on the timing of morphological landmarks of the SPB cycle. Note that cells continue to grow during a cell cycle block, and when they are released from their arrest, they typically progress quickly through the G1 phase of the cycle. In contrast, early G1 cells obtained by centrifugal elutriation are not arrested and must meet their growth requirements for Start. Hence, synchronous cultures obtained by centrifugal elutriation enable monitoring of early G1 events. To our knowledge, the SPB cycle has not been followed in elutriated synchronous cultures starting in early G1.

We used a strain harboring two fluorescently tagged SPB components, *Spc42p-GFP* and *Spc110p-tomato*, to visualize SPB duplication’s early steps. Spc42p is a central plaque component of the SPB (Bullitt *et al*. 1997), which Mps1p phosphorylates (Castillo *et al*. 2002). Mps1p also phosphorylates Spc110p, an inner plaque SPB component, which attaches γ-tubulin to the SPB (Friedman *et al*. 2001). Early G1 cells of this strain were obtained by elutriation and monitored at regular intervals for cell size and budding, and samples were fixed for SPB analysis by structured illumination microscopy (SIM) (Figure 3). The single SPB has a half-bridge attached, which doubles in length once SPB duplication commences (Li *et al*. 2006). Once the half-bridge elongates, proteins assemble at the distal end of the bridge to form a precursor to a new SPB called a satellite (Winey and Bloom 2012). Spc42p is one of several proteins deposited at the satellite location. This event is viewed as a cone-like structure flaring off the original SPB, denoted as ‘satellite-early’ in Figure 3. When most of the cells are unbudded, the prominent SPB feature seen is one SPB alone or one SPB with a small cone process, indicating early satellite body formation (Figure 3). Shortly before the size at which budding occurs in *uORF^+^-MPS1* cells, fully formed satellite processes are detected, denoted as ‘satellite-late’ in Figure 3. The appearance of duplicated, side-by-side SPBs is detected once the cells bud and pass through Start (Figure 3; (Byers and Goetsch 1974)). The final step, the separation of the SPBs, occurs during the anaphase of mitosis, which is consistent with our data showing this separation occurring near the end of the cell cycle (Figure 3). These data describe the morphology of early events of the SPB cycle in highly synchronous, elutriated newborn daughter cells, documenting the formation of early satellite structures soon after birth.

**FIGURE 3.**
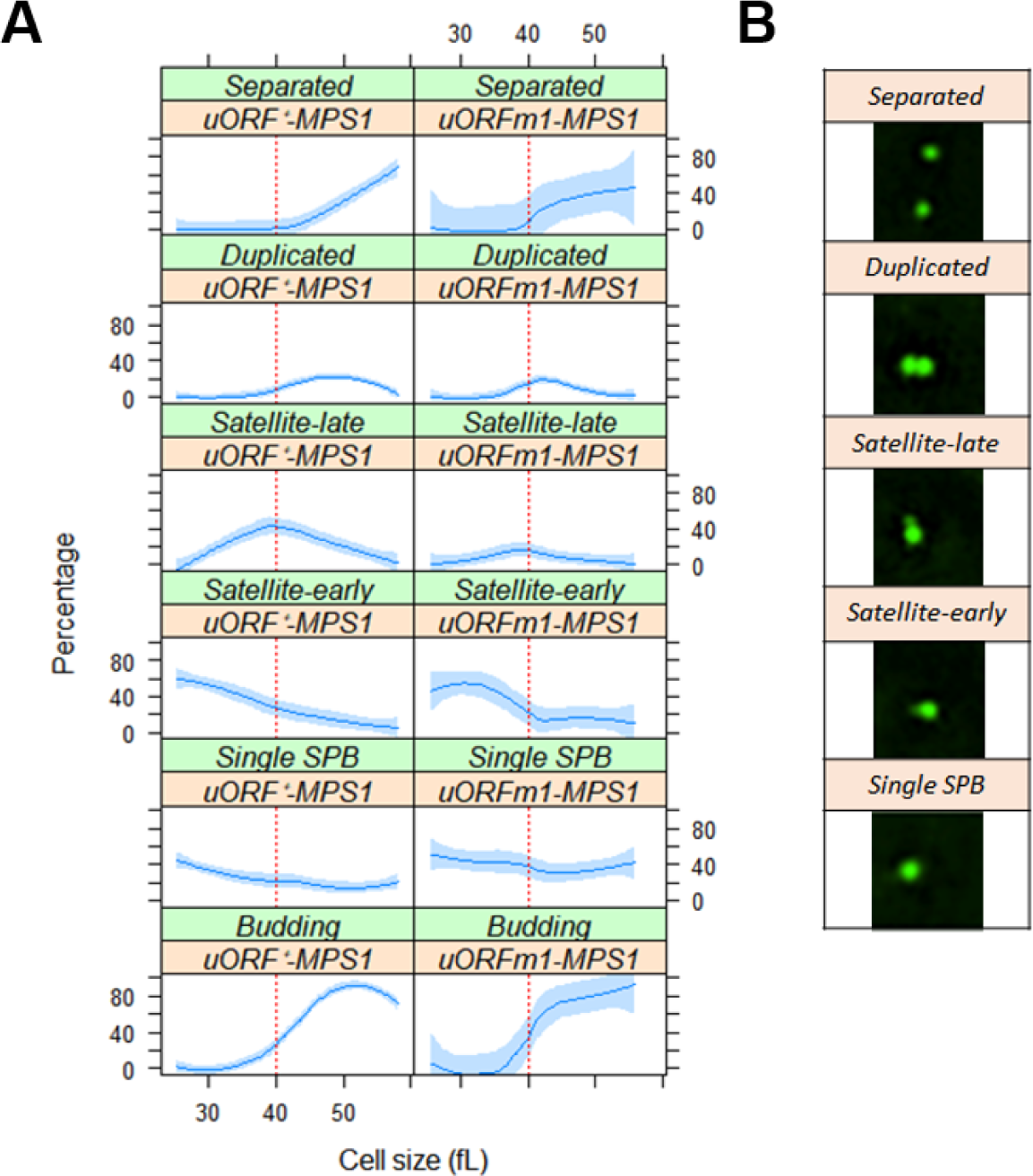
Reducing the translational efficiency of *MPS1* accelerates SPB duplication in synchronous, elutriated cells. **A**, Elutriated, early G1 daughter cells of the indicated *MPS1* uORF genotype carrying *SPC42-GFP* and *SPC110-tomato* alleles were monitored by SIM microscopy as they progressed synchronously in the cell cycle, in rich (YPD) medium. On the x-axis of each panel is cell size. On the y-axis is the percentage of cells with the indicated morphology, based on the fluorescence of SPB markers. Loess curves and the standard errors at a 0.95 level are shown. **B**, Representative SIM images of the SPB morphology, based on Spc42-GFP fluorescence, used to score the synchronous cells at various stages of the cell cycle. The values used to generate the graphs are in supplementary File1/sheet ‘fig3a’.

We introduced the *uORFm1-MPS1* allele in the *SPC42-GFP*, *SPC110-tomato* strain and measured the timing of the above event of the SPB cycle. Using cell size as a proxy for cell cycle position, the early steps of the SPB occur at the same time in the *uORFm1-MPS1* mutant and *uORF^+^-MPS1* cells. For example, the appearance of cells with satellite-late morphology peaks at 40 fL in both strains (Figure 3A). However, *uORFm1-MPS1* cells peak with SPB duplicated morphology soon after, at a size of ∼42 fL. In contrast, the *uORF^+^-MPS1* cells do so at a larger cell size (47-48 fL, compare the second-row panels in Figure 3A). The subsequent cell separation step is also accelerated in *uORFm1-MPS1* cells compared to *uORF^+^-MPS1* ones (Figure 3A, compare the two top panels). These results show that cells that do not have the *MPS1* uORF duplicate their SPB earlier, at a smaller size, consistent with an accelerated completion of Start.

### The accelerated Start of *MPS1* uORF mutants depends on Cln3p and Whi5p, but not on Cln1,2p

We then asked if *uORFm1-MPS1* functionally interacts with known regulators of Start. In yeast, two critical regulators of Start are the G1 cyclin Cln3p (promotes Start) and Whi5p (inhibits Start). Whi5p is a transcriptional repressor, analogous to the retinoblastoma (Rb) gene product of animal cells (Costanzo *et al*. 2004; de Bruin *et al*. 2004). Direct, localized phosphorylation of RNA PolII by the Cln3p-Cdc28p kinase complex triggers transcription of >100 genes at Start, including transcription of *CLN1* and *CLN2*, encoding G1/S cyclins (Kõivomägi *et al*. 2021). Removing Cln3p makes the cells bigger (Cross 1988; Nash *et al*. 1988), while removing Whip5 makes them smaller (Costanzo *et al*. 2004; de Bruin *et al*. 2004). Cells lacking Whi5p are among the smallest-sized mutants in yeast (Jorgensen *et al*. 2002). Introducing the *uORFm1-MPS1* allele in *whi5Δ* cells made them smaller (Figure 4A). On the other hand, mutating the *MPS1* uORF did not reduce the size of *cln3Δ* cells (Figure 4A). Lastly, Cln3p levels in the cell cycle did not change in *uORFm1-MPS1* mutants (Figure S3).

**FIGURE 4.**
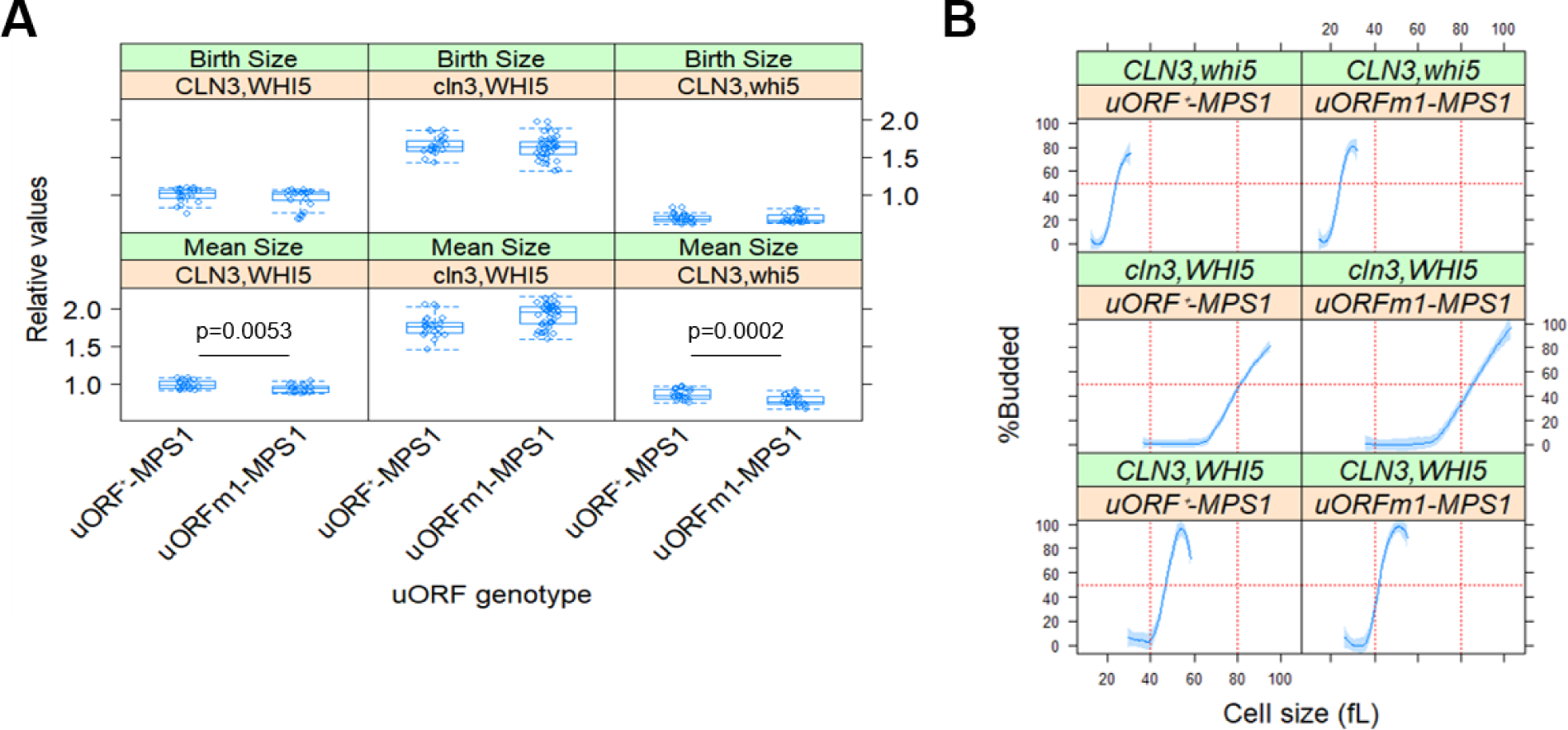
The smaller critical size of *MPS1* uORF mutants depends on the G1 cyclin Cln3p and the transcriptional repressor Whi5p. **A**, Box-plots showing the mean and birth size of populations of cells from multiple independent measurements from the indicated strains. The associated p-values from the non-parametric Wilcoxon rank-sum test between the two samples in each case are shown. **B**, Summary of cell cycle profiles of the indicated *MPS1* uORF genotype and strain background, done as in Figure 2. Loess curves and the standard errors at a 0.95 level are shown. The critical size is reduced by 9% (Wilcoxon rank sum test; p-value = 0.0005828) in *uORFm1-MPS1* cells, but there is no effect in *cln3Δ* and *whi5Δ* cells. The values used to generate the graphs are in supplementary File1/sheet ‘fig4a’ and sheet ‘fig4b’.

To better gauge the dependence of Mps1p on G1/S regulators, we measured the timing of Start in synchronous, elutriated cultures. The critical size threshold for Start is greatly increased in cells lacking Cln3p (at 80 fL, compared to 46 fL for wild type cells; Figure 4B), and it was not affected at all in double, *uORFm1-MPS1, cln3Δ* mutants (Figure 4B). Likewise, despite their smaller size (Figure 4A), *uORFm1-MPS1, whi5Δ* mutants did not have a reduced critical size at START (Figure 4B). A caveat in the analysis of the critical size of *whi5Δ* cells is that these mutants have such a small critical size, so a further reduction by de-repressing translation of *MPS1* may be difficult to impose. These results suggest that Cln3p, and possibly Whi5p, are needed for Mps1p to accelerate Start when the *MPS1* uORF is mutated.

We also measured Start kinetics in cells lacking both the Cln1p and Cln2p G1 cyclins. Double *cln1,2Δ* mutants have a much larger critical size (at ∼71 fL, compared to 46 fL for wild type cells; Figure 5A). De-repressing translation of *MPS1*, in *uORFm1-MPS1, cln1,2Δ* cells, significantly reduced critical size (∼60 fL; Figure 5A). We also noticed that loss of Cln1,2p dampened the cell cycle-dependent oscillation of Mps1p (Figure 5B, C), but it was restored in *uORFm1-MPS1, cln1,2Δ* cells (Figure 5B, C). Nonetheless, despite the smaller amplitude in the changing levels of Mps1 in the cell cycle in *cln1,2Δ* cells, these results suggest that de-repressing translation of *MPS1* accelerates Start mostly independently of Cln1,2p.

**FIGURE 5.**
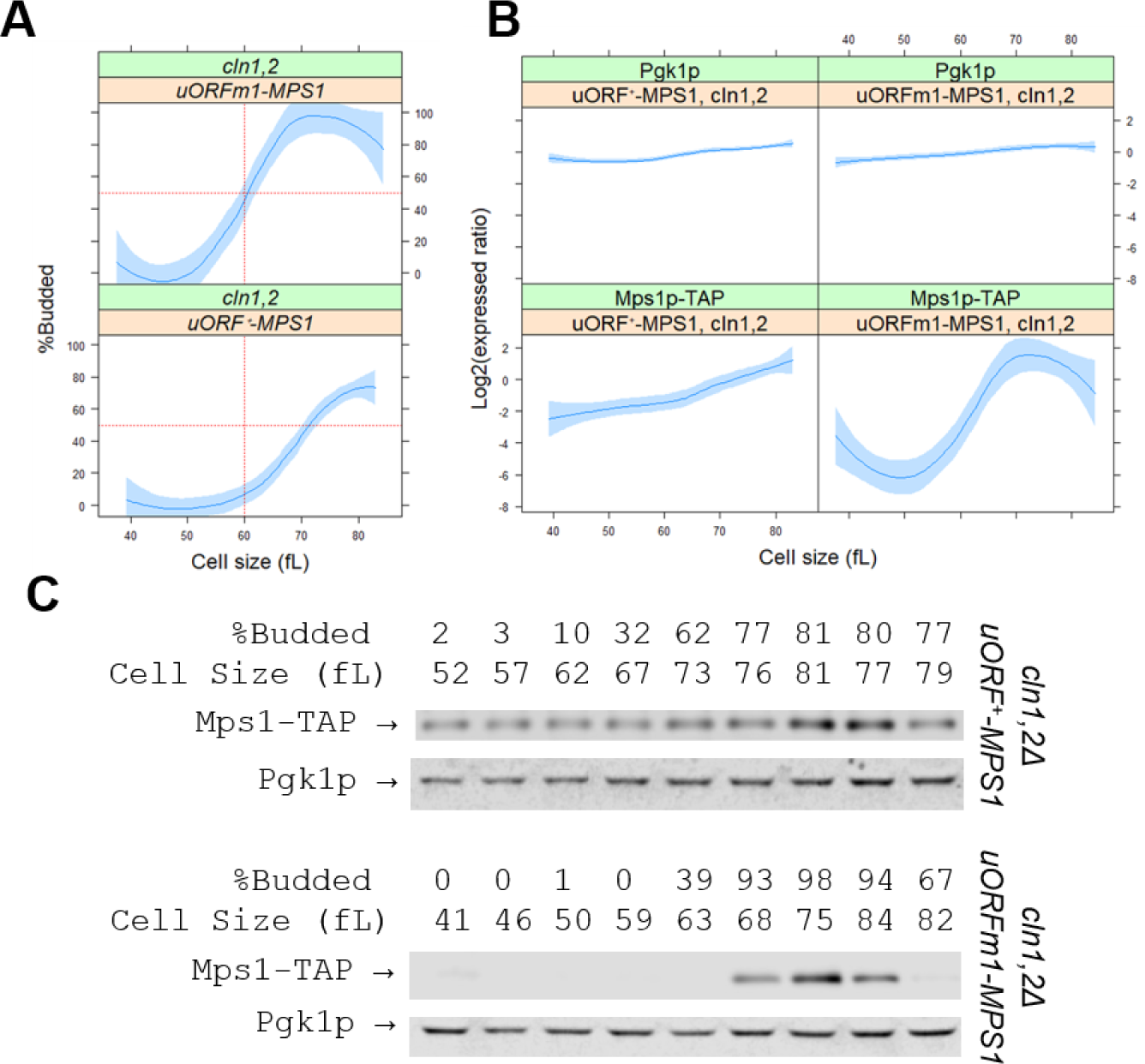
Translational de-repression of Mps1p accelerates Start in cells lacking Cln1p and Cln2p. **A,** Summary of cell cycle profiles of double *cln1,2Δ* mutants, of the indicated *MPS1* uORF genotype background, done as in Figures 2&3. Loess curves and the standard errors at a 0.95 level are shown. The critical size is reduced by at least 10 fL in *uORFm1-MPS1* cells. The values used to generate the graphs are in supplementary File1/sheet ‘fig5a’. **B**, The amplitude of the Mps1p-TAP oscillation in the cell cycle is smaller in *cln1,2Δ* mutants, but it is restored when mutating the *MPS1* uORF. The abundance of Mps1p-TAP was monitored as in Figure 2. **C**, Representative immunoblots, along with the percentage of budded cells (%B) and the cell size (in fL), for each of the samples. All the immunoblots for this figure are in supplementary File4, while the values used to generate the graphs are in supplementary File1/sheet ‘fig5a’ and sheet ‘fig5b’.

Since Cdk is known to phosphorylate Mps1p at T29 (Jaspersen *et al*. 2004), it is possible that phosphorylation of Mps1p by G1/Cdk somehow promotes the G1/S transition. To test this idea, we measured the critical size of the non-phosphorylatable *MPS1-T29A* mutant. There was no difference in the critical size of *MPS1-T29A* mutant cells and their otherwise wild type counterparts (Figure S4). These data suggest that the Cdk-dependent phosphorylation of Mps1 does not significantly affect the timing of Start. Instead, Mps1p may promote Start by phosphorylating some G1/S regulators. We tested this possibility in vitro, asking if Mps1p could phosphorylate Whi5p or Sic1p (Figure S5). Sic1p is an inhibitor of B-type cyclin/Cdk complexes, whose phosphorylation triggers its degradation, enabling DNA replication to commence (Venta *et al*. 2020). Indeed, in vitro, Mps1p phosphorylated both Sic1p and Whi5p (Figure S5A). However, Mps1p’s kinase activity towards Sic1p and Whi5p was lower than that of Cln2/Cdk and Clb5p/Cdk towards these substrates (Figure S5A). We then examined the ability of Mps1p to phosphorylate Whi5p and Sic1p compared to other known Mps1p substrates with roles in SPB duplication (Cdc31p and Spc29p) or the spindle assembly checkpoint (Ndc80p). The Mps1p activity toward Sic1p and Whi5p was comparable to its activity against Cdc31p but 2-3-fold lower than its activity against Ndc80p and Spc29p (Figure S5B). Furthermore, Mps1p could target Sic1p even when all known Cdk phosphorylation sites on Sic1p were mutated to alanine (Sic1p-AP; see Figure S5B), presumably at other non-Cdk sites. Although it is currently unclear how Mps1p exercises its role in the timing of Start in cells, our in vitro data argue that Mps1p could potentially phosphorylate Start regulators (Figure S5). In such a scenario, Mps1p role may be secondary to Cdk or other kinases, helping to bring about this critical cellular transition, possibly through its broad specificity as a kinase. We note that, in a perhaps analogous manner, Mps1p ensures that the spindle assembly checkpoint is established, participating in the phosphorylation of outer kinetochore proteins following the action of the Aurora-B kinase (Liu and Winey 2012).

## Conclusions

The role of centrosomes in cell division is as old as the chromosome theory of heredity, with centrosome amplification a common feature of cancers (Boveri 2008). Normally, centrosome and SPB duplication are entrained with the master cell cycle switch governed by the activity of cyclin-dependent kinase (Cdk) (Jaspersen and Winey 2004; Nigg and Raff 2009). However, in some cases, they can duplicate independently of cell-cycle progression (Nigg and Raff 2009). In animals, autonomous oscillations of the polo-like kinase Plk4 can drive centrosome duplication independently of Cdk (Aydogan *et al*. 2020). Intriguingly, centriole growth in the cell cycles of fly embryos appears to follow a homeostatic ‘sizer-like’ model, actively monitoring and adjusting their growth until they reach the proper size (Aydogan *et al*. 2018). An analogous mechanism has not been shown for the budding yeast SPB. However, the same principle operates during the G1 phase of daughter cells to control Cdk activity before cells commit at Start to initiate budding and DNA replication (Hartwell and Unger 1977; Di Talia *et al*. 2007). Our results argue that through translational control of *MPS1*, growth and protein synthesis also directly trigger SPB duplication, expanding the range of growth inputs the cell uses to control the timing of Start (Figure 6). Hence, monitoring the growth and duplication of the SPB/centrosome may contribute to the overall coupling of cell growth with division in proliferating cells.

**Figure 6.**
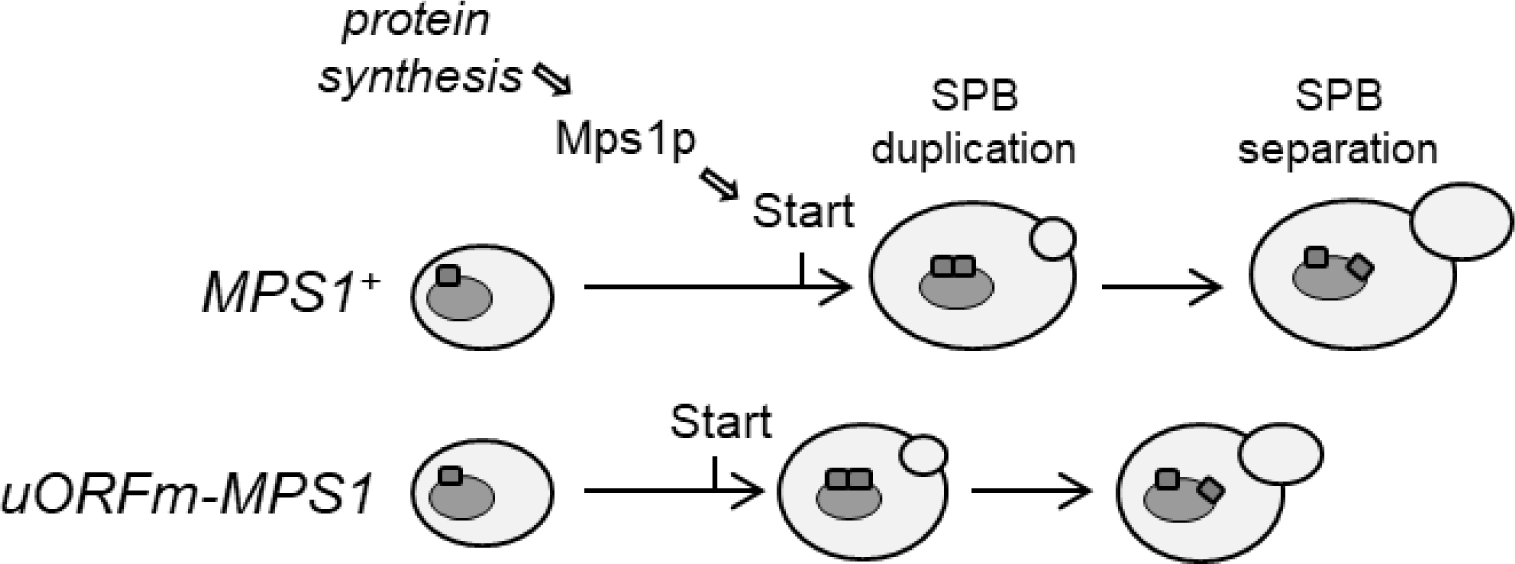
Schematic overview of our findings, connecting protein synthesis to Start and SPB duplication processes via translational regulation of *MPS1*.

## DATA AVAILABILITY

Strains and plasmids are available upon request. The authors affirm that all data necessary for confirming the conclusions of the article are present within the article, figures, and tables.

## ACKNOWLEDGEMENTS

We thank the MCB Light Microscopy Imaging Facility at the University of California - Davis. We also thank Dr. Mart Loog (University of Tartu, Estonia) for the in vitro kinase assays.

---

**Figure S1.**
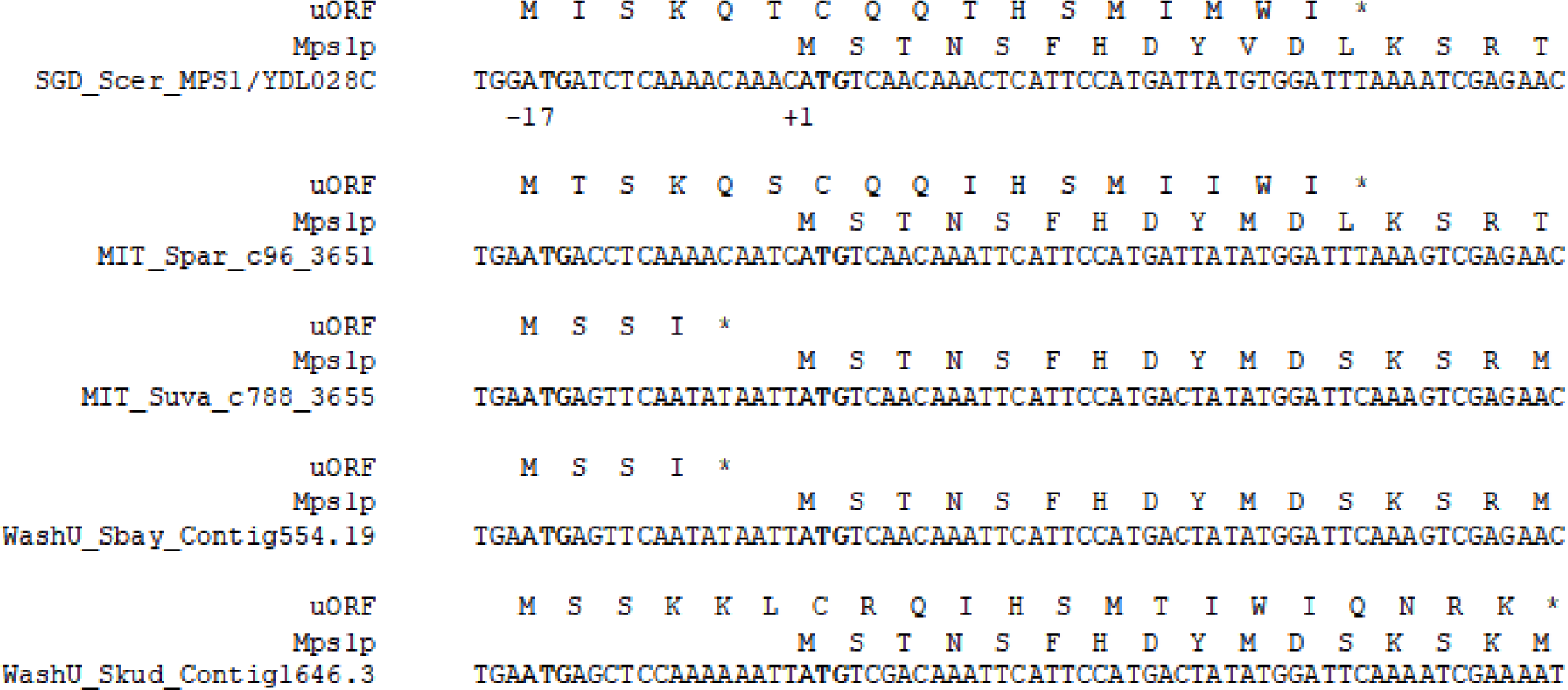
The initiation position of a uORF in the *MPS1* transcript, but not the length or the sequence of the uORF-encoded peptide, is conserved in the *Saccharomyces* genus. The genomic DNA sequence for *S. cerevisiae* (top) and for the related species are shown (as listed on the Saccharomyces Genome Database (SGD) fungal sequence alignments (Cherry *et al*. 2012)). Above each DNA sequence, the sequences of the uORF-encoded peptides and the amino-terminal sequence of Mps1p are shown.

**Figure S2.**
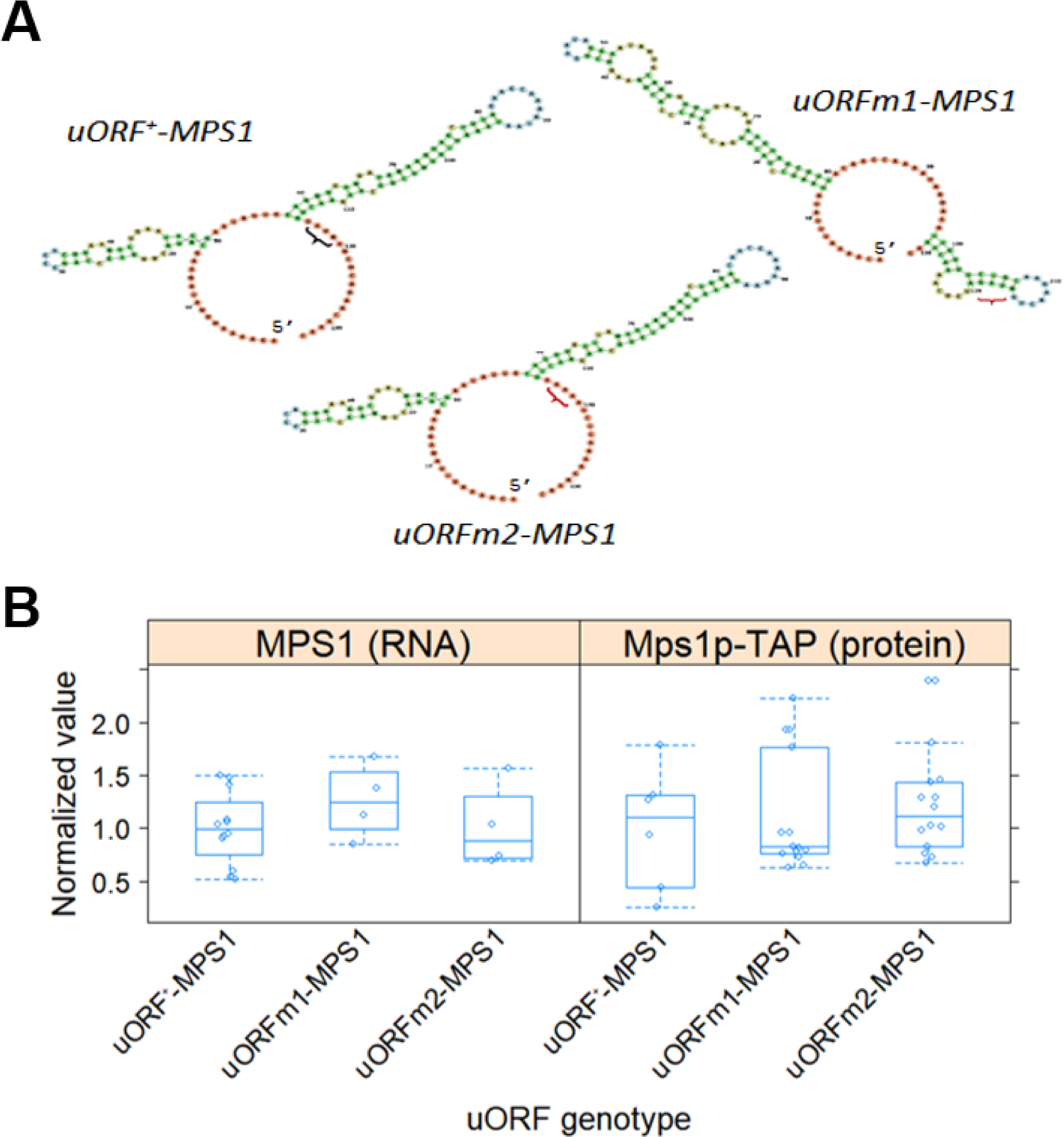
Mutating the *MPS1* uORF does not significantly change the steady-state abundance of Mps1p in asynchronous cells. **A**, Predicted RNA structure of the *MPS1* 5’-leader in each of the strains shown. The predicted structures were the minimum free energy (MFE) ones calculated with the Vienna RNA Websuite (Gruber *et al*. 2008). The brackets indicate the position of the start codon of the uORF in each case. **B**, Mutating the *MPS1* uORF does not significantly change the steady-state abundance of Mps1p in asynchronous cells. The box-plots show the levels of *MPS1* mRNA (left panel) and Mps1p protein (right panel) from multiple independent experiments, measured as described in Materials and Methods. The values were normalized against both Pgk1p levels in each sample and total protein, using the Mini-PROTEAN® TGX Stain-Free™ Gels, as described in Materials and Methods. All the immunoblots for this figure are in supplementary File3, while the values used to generate the graphs are in supplementary File1/sheet ‘figs2b’.

**Figure S3.**
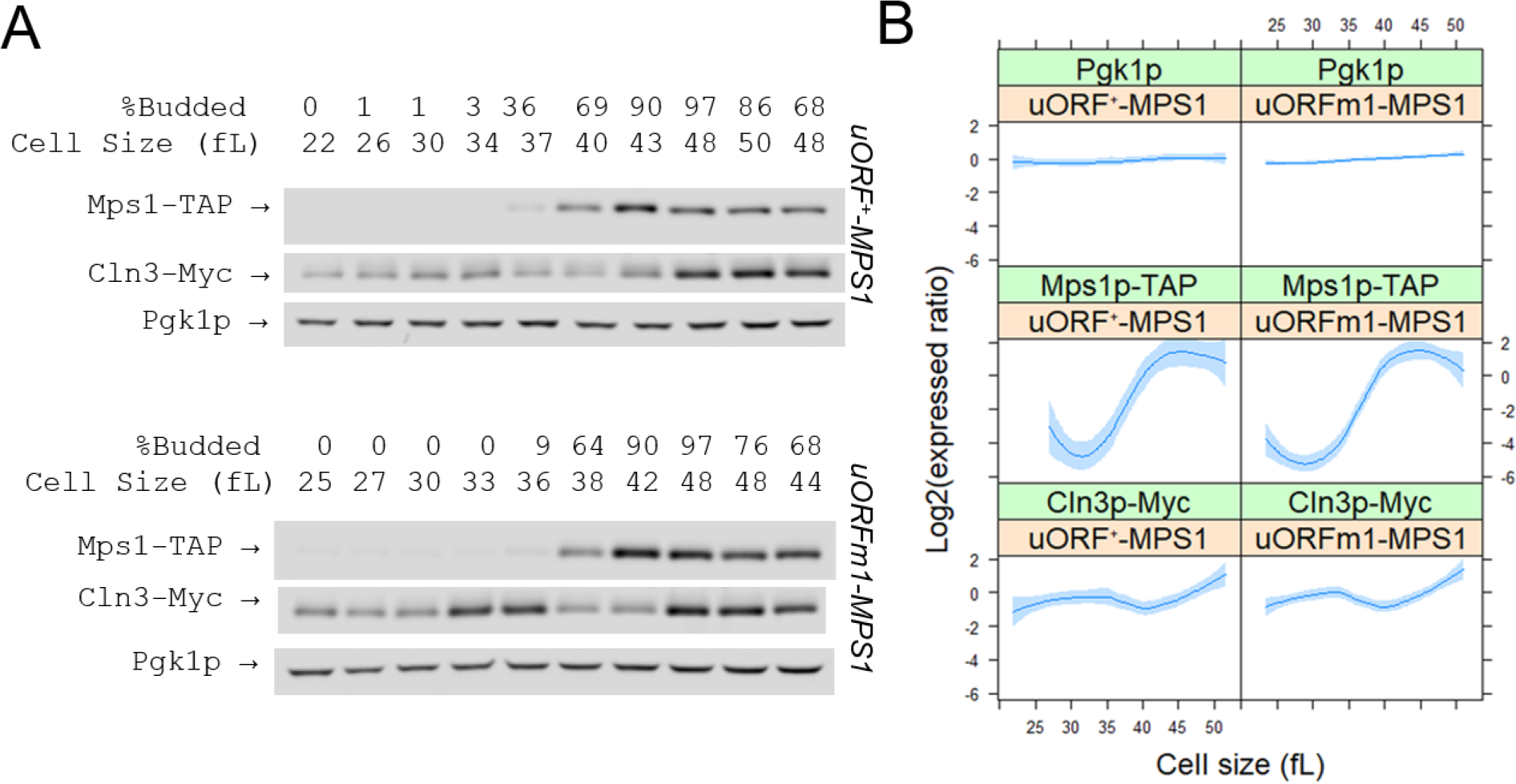
Translational de-repression of Mps1p does not change Cln3p levels in the cell cycle. The abundance of the indicated proteins was monitored in wild type and *uORFm-MPS1* cells, constructed as described in Materials and Methods. Samples were collected by elutriation in a rich, undefined medium (YPD) and allowed to progress synchronously in the cell cycle. Experiment-matched loading controls (measuring Pgk1p abundance) were also quantified and shown in parallel. **A**, Representative immunoblots, along with the percentage of budded cells (%B) and the cell size (in fL), for each of the samples. **B**, From at least three independent experiments in each case, the Mps1p-TAP, Cln3-Myc, and Pgk1p signal intensities were quantified as described in Materials and Methods. The Log2(expressed ratios) values are on the y-axis, and cell size values are on the x-axis. Loess curves and the standard errors at a 0.95 level are shown. All the immunoblots for this figure are in supplementary File5, while the values used to generate the graphs are in supplementary File1/sheet ‘figs3b’.

**Figure S4.**
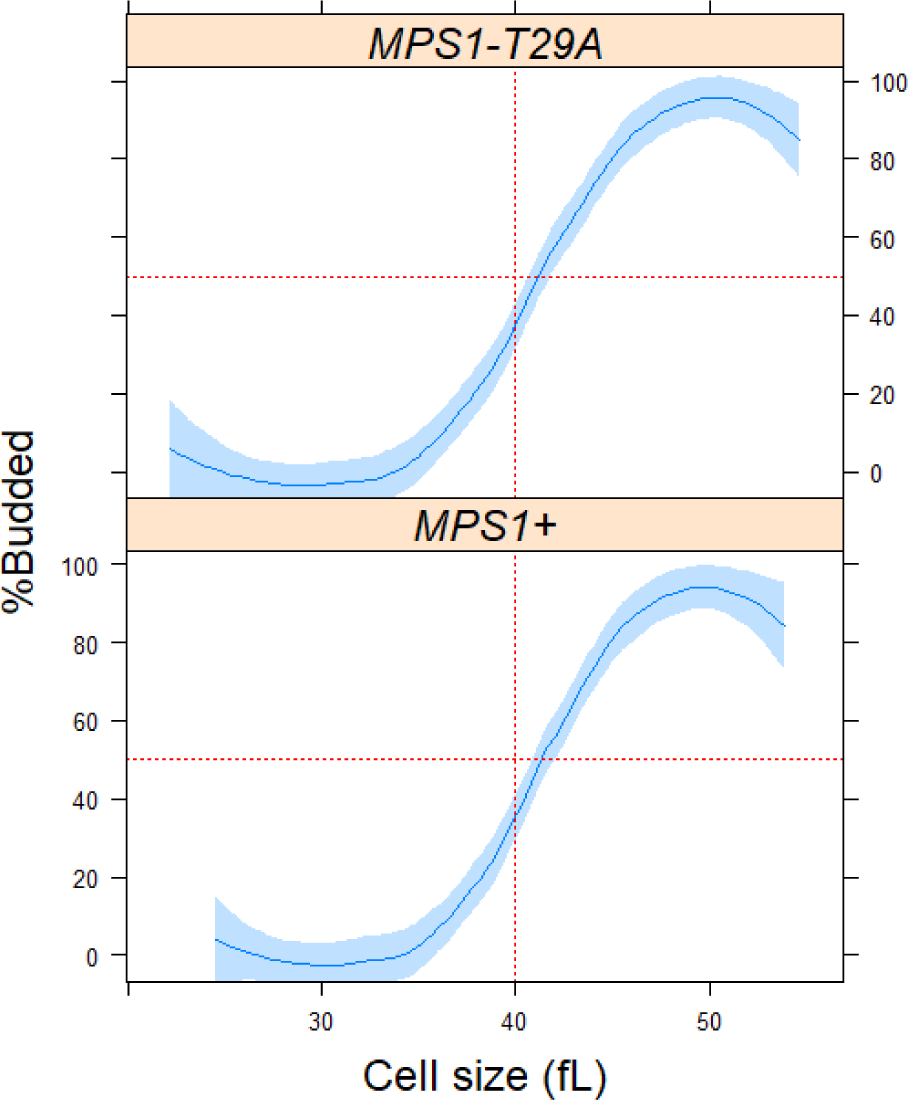
The timing of Start is not affected by introducing to Mps1p the T29A substitution that cannot be phosphorylated by Cdk. Summary of cell cycle profiles of the indicated strains, done as in Figure 2. Loess curves and the standard errors at a 0.95 level are shown. The values used to generate the graphs are in supplementary File1/sheet ‘figs4’.

**Figure S5.**
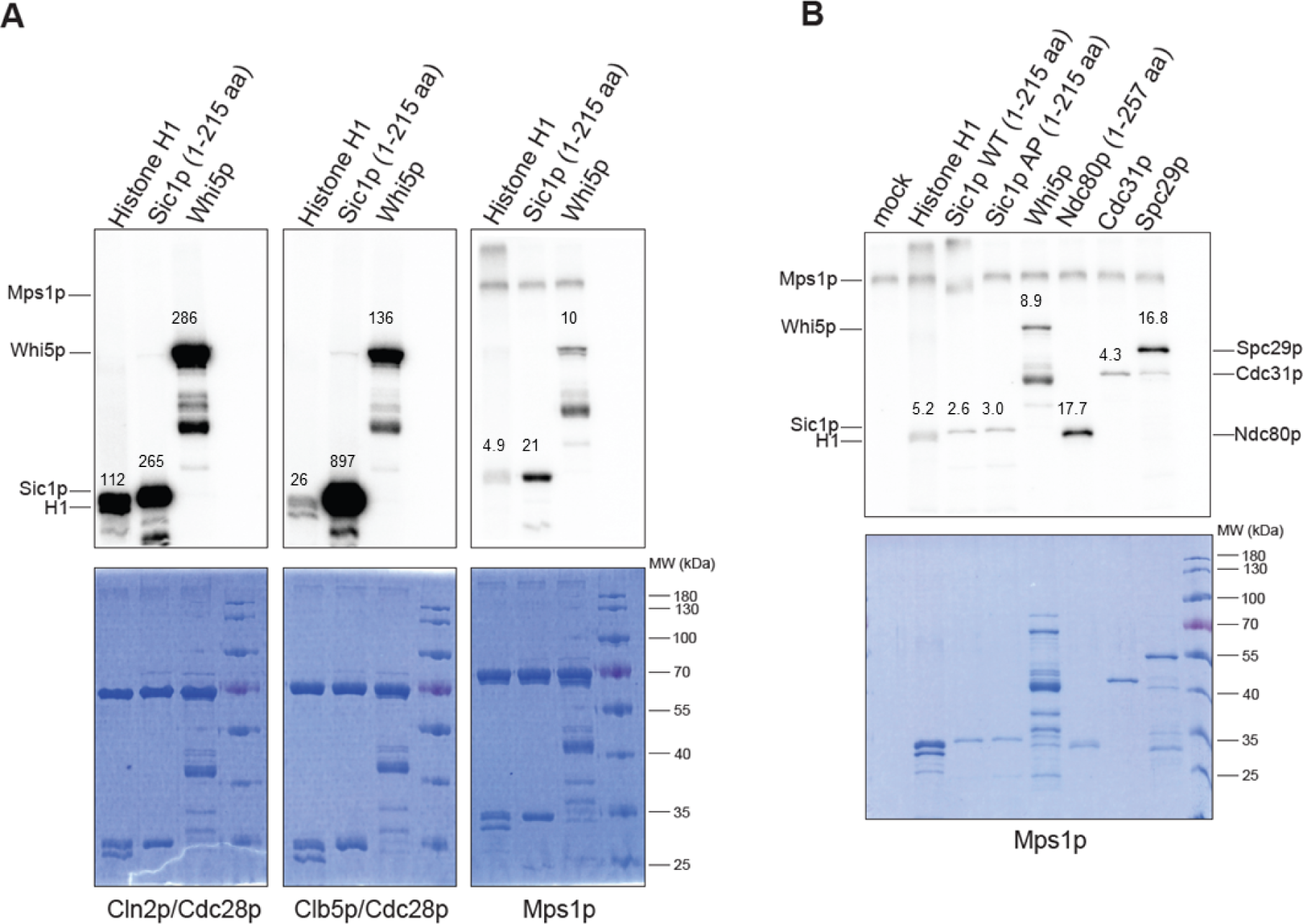
Mps1p can phosphorylate Whi5p and Sic1p in vitro. Autoradiograms of in vitro kinase reactions, displaying the incorporation of [^32^P]-γ-ATP into the indicated substrates resolved on SDS-PAGE gels are shown top. The Coomassie-stained gels are shown below the autoradiograms. **A**, The kinases shown at the bottom were incubated with the substrates shown at the top for 16m, as indicated. **B**, Mps1p was incubated with each substrate shown at the top for 16m. The mock samples contain Mps1p alone. The numbers above the bands corresponding to each of the protein substrates are the relative quantification of the signal on each gel.

