## Supplementary figures and images for "Translational control of *MPS1* links protein synthesis with the initiation of cell division and spindle pole body duplication in *Saccharomyces cerevisiae*"

### File2

#### supplementary File2

##### Mps1p levels during synchronous progression through the cell cycle.

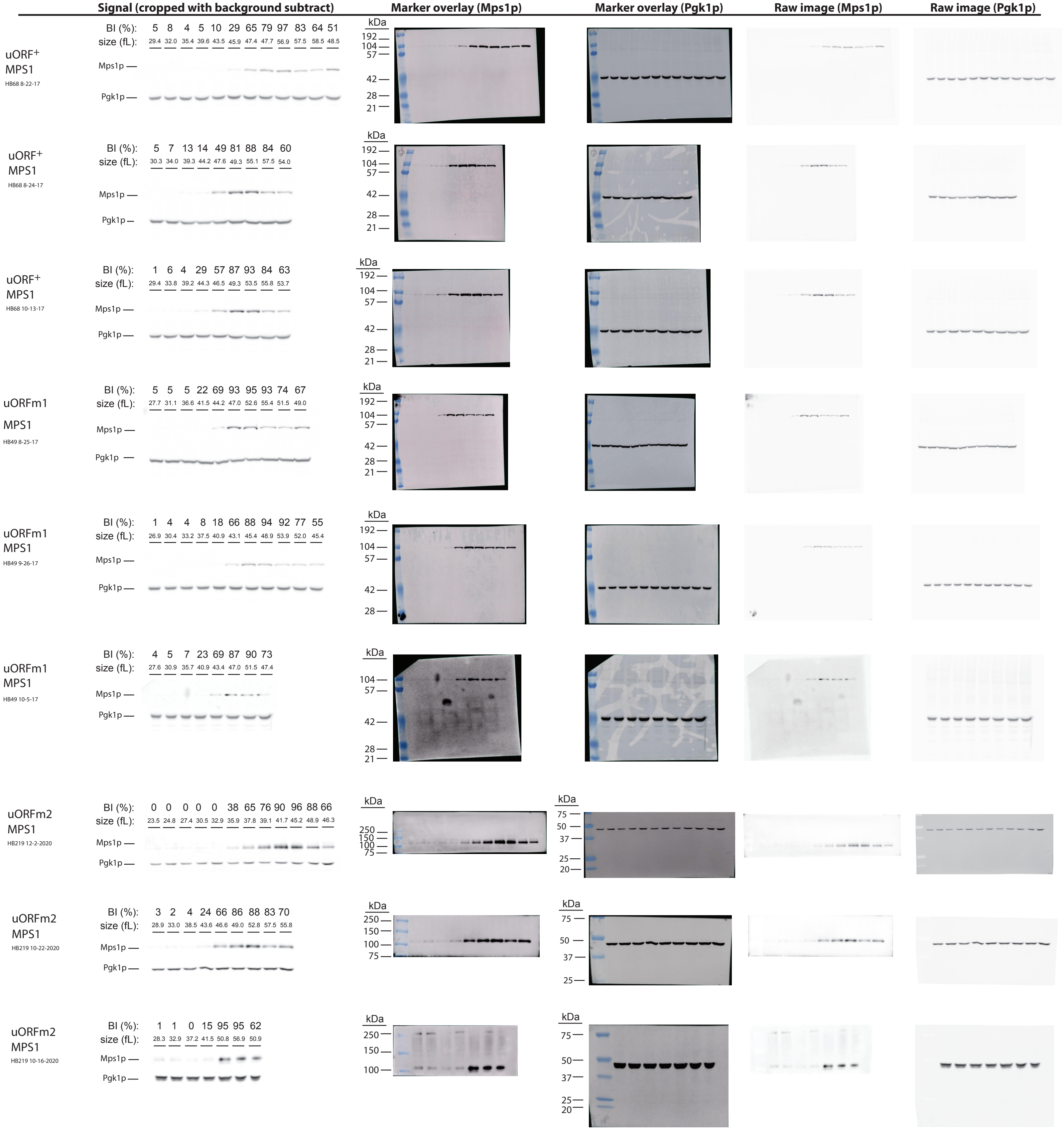

### File5

#### supplementary File5

##### uORFm1 MPS1-TAP CLN3-Myc

### HB264 3-8-2021

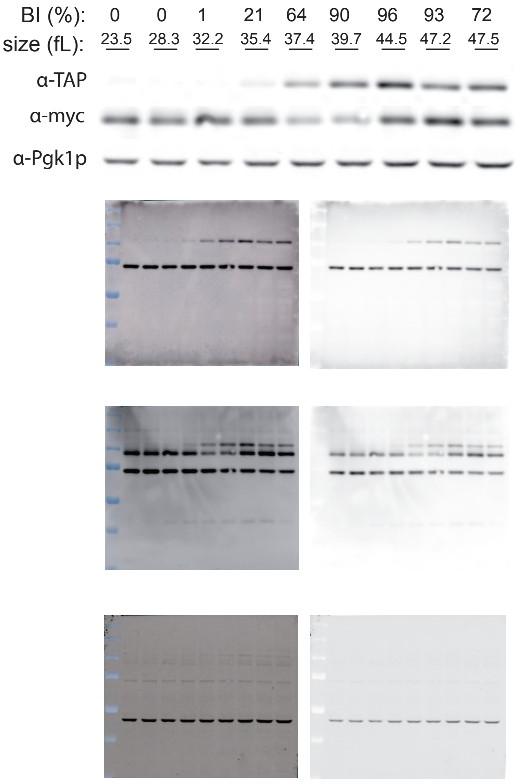


### HB264 3-10-2021

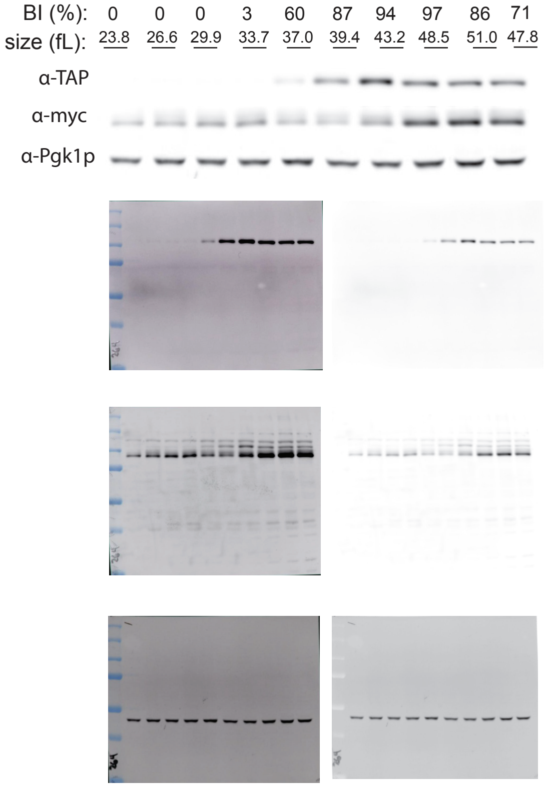


### HB264 2-11-2021

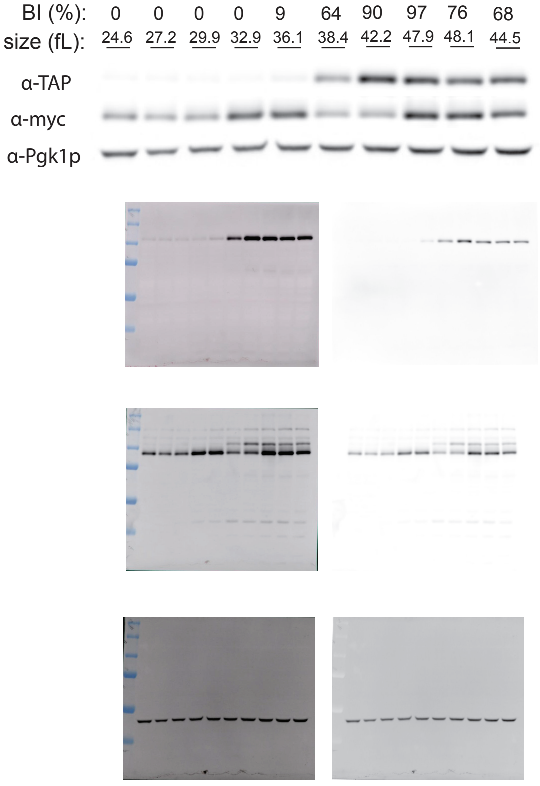


##### uORF⁺ MPS1-TAP CLN3-Myc

### HB270 3-10-2021

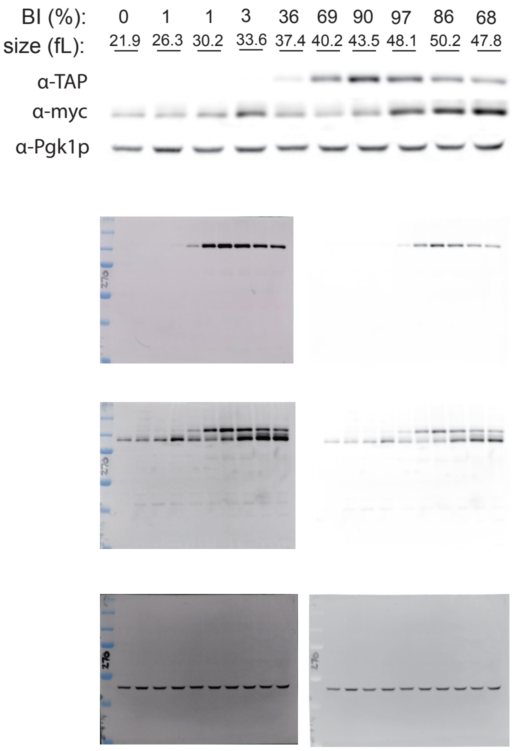


### HB270 2-23-2021

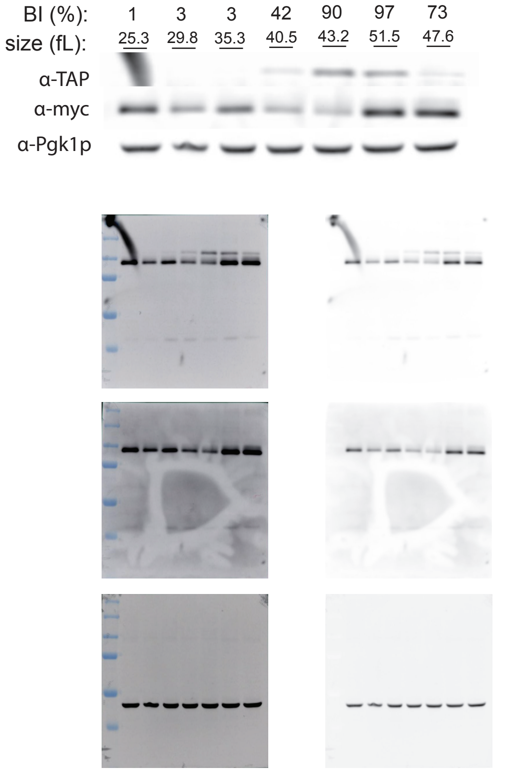


### HB270 2-26-2021

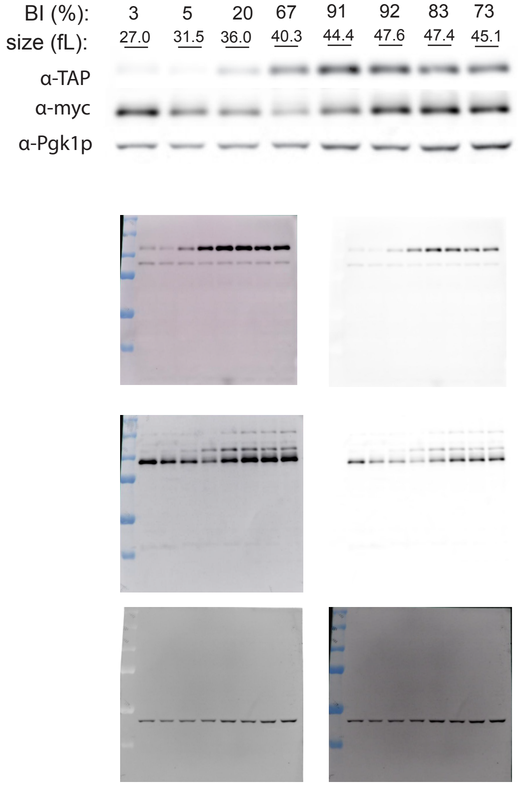
