## Supplementary material for "Translational control of *MPS1* links protein synthesis with the initiation of cell division and spindle pole body duplication in *Saccharomyces cerevisiae*": File3

### supplementary File3

#### Mps1p levels in batch cultures.

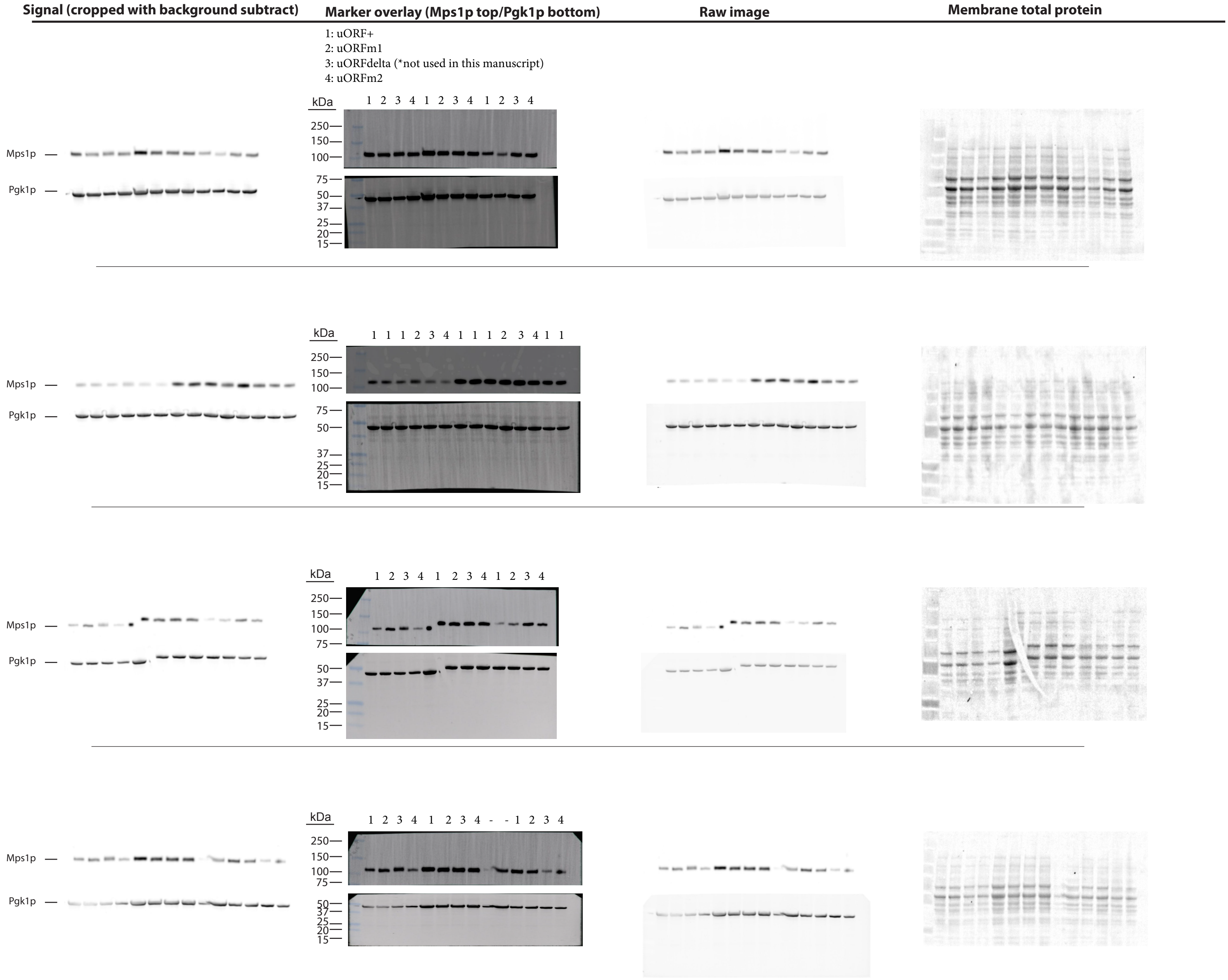


1: uORF+
2: uORFm1
3: uORFdelta (*not used in this manuscript)
4: uORFm2
