## Supplementary material for "Translational control of *MPS1* links protein synthesis with the initiation of cell division and spindle pole body duplication in *Saccharomyces cerevisiae*": File4

### supplementary File4

Immunoblots from uORF+ and uORFm1 cells lacking CLN1, CLN2.

#### *cln1Δ cln2Δ uORFm1-MPS1-TAP*

### HB281 4-8-2021

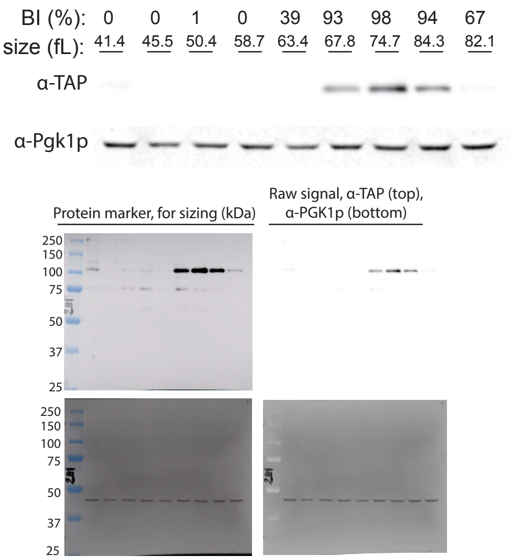


### HB281 4-15-2021

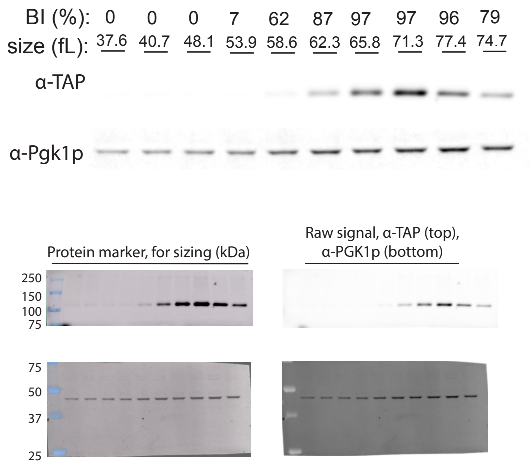


### HB281 6-16-2021

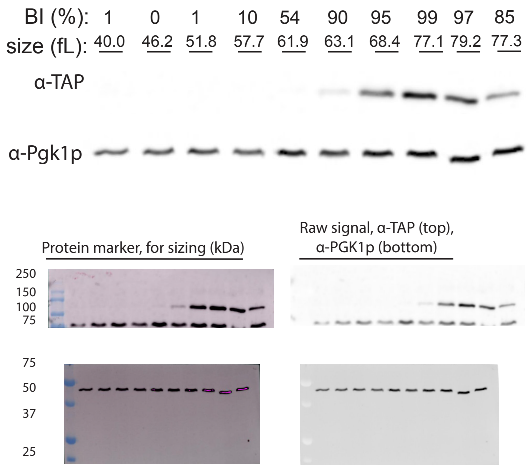


#### *cln1Δ cln2Δ uORF⁺-MPS1-TAP*

### HB278 4-28-2021

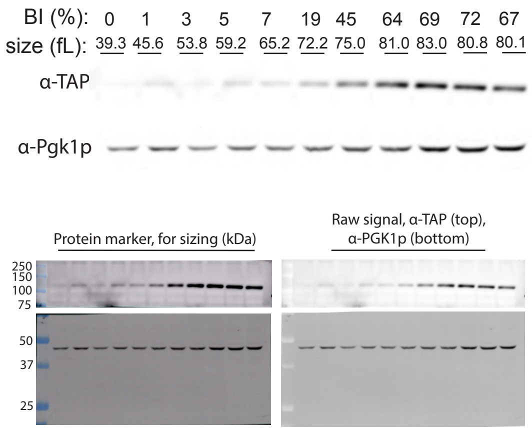


### HB278 4-9-2021

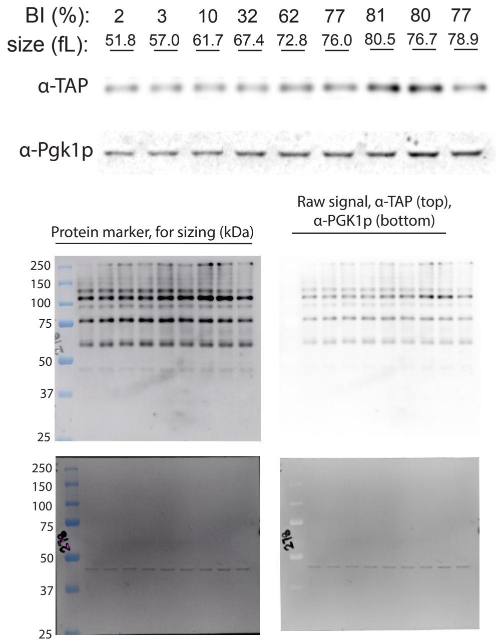


### HB278 4-13-2021

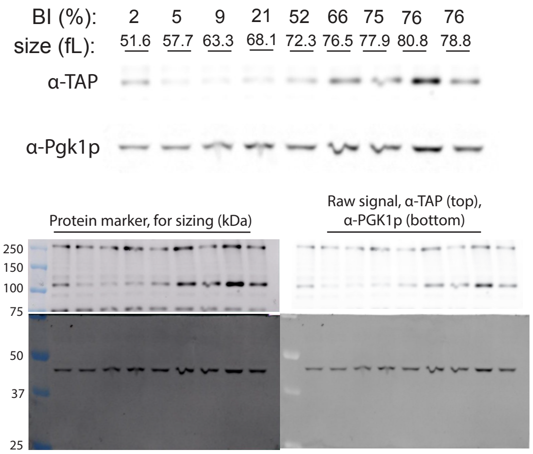


### HB280 4-30-2021

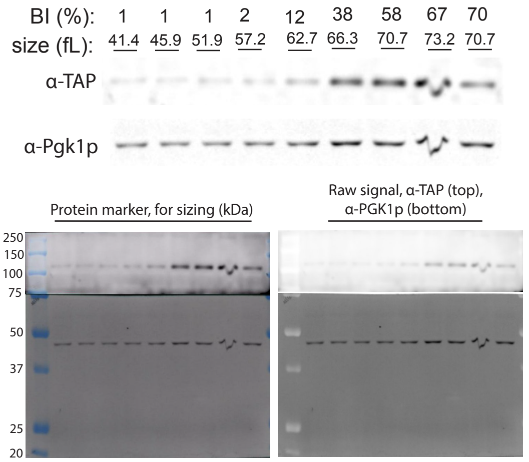
